# Ferroptosis links α-synuclein pathology across brain and skeletal muscle in Parkinson’s disease

**DOI:** 10.64898/2026.04.08.717156

**Authors:** Krishna Singh Bisht, Jaydeep Sharma, Naman Kharbanda, Ankit Biswas, Sandhini Saha, Sam Jacob Mathew, Tushar Kanti Maiti

## Abstract

Parkinson’s disease (PD) is increasingly recognized as a multisystem disorder, yet the mechanisms linking neurodegeneration with muscle dysfunction remain largely unknown. In this study, using an A53T α-synuclein (αSyn) transgenic mouse model, we demonstrate coordinated pathological changes across the brain-muscle organs characterized by systemic inflammation, iron accumulation, oxidative stress, and ferroptosis-associated lipid peroxidation. Our quantitative proteomics data revealed dysregulated iron metabolism and ferroptosis in the brain and skeletal muscle. Biochemical validation confirmed increased expression of Transferrin receptor 1 (TFRC), elevated lipid peroxidation, and suppression of antioxidant defenses, including SLC7A11 and GPX4, indicating enhanced ferroptotic susceptibility. Cell-surface proteomics and biophysical assays further revealed that pathological αSyn directly interacts with TFRC, promoting iron accumulation and ferroptosis-associated oxidative damage in neuronal and muscle cells. Together, our findings identify ferroptosis as a shared pathological mechanism across the brain and muscle, mediated by the αSyn-TFRC interaction, thus linking neurodegeneration and peripheral muscle pathology in PD.

## Introduction

Parkinson’s Disease (PD) is a multifactorial progressive neurodegenerative disease affecting millions worldwide. PD predominantly affects the elderly population above the age of 60 years, but cases of early-onset PD are on the rise, affecting people below the age of 30[1]. The accumulation of proteinaceous aggregates called Lewy bodies (LBs) in the substantia nigra pars compacta (SNpc) region is a major pathological hallmark of PD[2,3]. Misfolded α-synuclein is a major component of LBs[3]. PD is associated with the degeneration of dopaminergic neurons in the SNpc, a brain region responsible for motor coordination. Therefore, the major symptoms of the disease involve akinesia, bradykinesia, tremors, and postural instability. However, long before motor symptoms emerge, individuals often experience cognitive impairments, anosmia, constipation, sleep disturbances, and autonomic dysfunction[4]. PD is now increasingly recognized as a multisystem disease involving peripheral organs[5]. However, the molecular mechanisms that mechanistically couple central neurodegeneration to peripheral dysfunction remain poorly defined.

α-Synuclein (αSyn) is an unstructured protein that associates with synaptic vesicle membranes and participates in synaptic vesicle trafficking, SNARE complex assembly, and neurotransmitter release, and is a central driver of PD pathogenesis[6,7]. *SNCA* gene, encoding αSyn, mutations, namely A53T, A30P, H50Q, E83Q, A18T, G51D, and E46K, as well as gene duplication and triplication, are known to induce aberrant αSyn aggregation and toxicity[2,8,9]. Although previously considered confined to the brain, αSyn aggregation is increasingly recognized as systemically present. Several studies have shown that the αSyn pathology is present in the enteric nervous system, cardiac tissue, skeletal muscle, kidneys, liver, and even cutaneous nerves[10–17]. These observations suggest that αSyn pathology may also begin at peripheral sites and ascend to the brain via specific neural pathways[18].

Along with intracellular αSyn accumulation, extracellular αSyn can engage cell-surface receptors, such as TLR2, RAGE, LAG3, etc., and propagate its spreading and pathological signaling across cell types. These receptor-mediated interactions have been implicated in αSyn endocytosis, neuroinflammation, synaptic dysfunction, and the spread of pathology[19–23]. However, the surface molecules linking αSyn to metabolic and redox dysfunction, particularly across multiple organs, remain unclear. Oxidative stress and iron dyshomeostasis are key contributors to PD pathology[24]. Iron deposition in the SNpc of PD patient brains has been associated with dopaminergic neuron loss via a process called ferroptosis[25–27]. Furthermore, inhibition of ferroptosis can alleviate motor deficits and neuronal loss in MPTP-induced PD mice[28]. Ferroptosis is an iron-dependent form of regulated cell death initiated by abnormal iron metabolism, which promotes the generation of reactive oxygen species and severe lipid peroxidation, leading to oxidative stress and cell death[29]. Ferroptosis is driven by lipid peroxide accumulation, causing membrane damage and failure of antioxidant defenses controlled by NRF2 and its downstream targets, including SLC7A11 (Solute Carrier Family 7 Member 11) and GPX4 (Glutathione peroxidase 4)[30,31]. Iron accumulation and αSyn pathology have been closely associated with colocalization in the midbrain[32]. αSyn also binds ferric and ferrous iron, promoting αSyn aggregation[33]. αSyn mRNA contains an iron response element (IRE) within its 5′-untranslated region (UTR). Thus, iron also participates in post-transcriptional regulation of αSyn[34,35]. Conversely, αSyn can also modulate iron homeostasis as its overexpression in rat primary midbrain neurons results in iron overload[36], and the ferrireductase activity of αSyn can increase intracellular ferrous iron content as previously shown in SH-SY5Y cells[37]. However, the intricate mechanisms underlying αSyn and iron-induced toxicity remain unclear. While ferroptosis has been increasingly linked to neuronal vulnerability in PD, whether it represents a shared mechanism across central and peripheral tissues remains unknown.

Skeletal muscle dysfunction is a clinically significant feature of PD, contributing to motor impairments, accelerated disease progression, and reduced quality of life[38]. PD is closely associated with sarcopenia or loss of skeletal muscle mass and function, leading to impaired posture, movement, and locomotor capability[39]. Further PD-associated sarcopenia increases the incidence of falls and related muscle injury, making routine daily activities risk-prone[40,41]. The presence of αSyn in skeletal muscle suggests that peripheral tissues may be directly affected in PD[15]. Skeletal muscle, in addition to its motor functions, also acts as an endocrine organ that communicates with the brain through myokines and metabolites[42]. Thus, in PD, disrupted muscle function and altered signaling may both result from and contribute to ongoing neurodegeneration[41,43]. However, it remains unclear whether common mechanisms, such as iron dysregulation and ferroptosis, can drive degeneration in both brain and muscle, or how these processes may be coordinated systemically. The reports on the role of αSyn in the brain are abundant, but studies linking it to the skeletal muscle in PD are limited.

In this study, using a transgenic mouse model of PD expressing human A53T αSyn, we investigate αSyn-associated pathology across the brain and skeletal muscle. Aged A53T mice exhibited PD-associated motor impairment along with reduced body and muscle weight. These mice showed increased αSyn accumulation in both the brain and skeletal muscle, along with neurodegeneration and neuromuscular junction (NMJ) defects. Through integrated multi-proteomics and biochemical analyses in A53T α-synuclein mouse model, and cell-surface proteomics in neuronal and muscle cells, we identify a novel αSyn–TFRC–ferroptosis axis that operates across both the brain and skeletal muscle. Mechanistically, αSyn directly interacts with TFRC to enhance iron uptake, suppress antioxidant defenses, and promote lipid peroxidation. Additionally, plasma proteomic profiling in A53T mice reveals a systemic ferroptosis-associated signature that links brain and skeletal muscle pathology. Together, these findings highlight ferroptosis as a mechanism linking brain–muscle pathology and expanding current understanding of PD as a systemic disorder driven by αSyn-mediated iron dysregulation.

## Results

### 1. A53T mice exhibit αSyn-induced Parkinson’s Disease symptoms in the brain and muscle

Extensive PD studies have examined the role of αSyn in the brain, but investigations connecting the brain and skeletal muscle pathology remain limited. Using a transgenic mouse model of PD expressing human A53T αSyn, we investigate αSyn aggregation-associated pathology in both brain and skeletal muscle. For this purpose, we used 18-month-old hemizygous *A53T ^(+/-)^* mice and their age-matched littermate non-transgenic (control) mice. The A53T mice showed decreased body weight compared to controls (Figure 1A). On the rotarod, A53T mice showed decreased latency to fall, whereas they took a longer duration to descend from the pole than controls (Figure 1B, C), thus indicating motor control impairment in the A53T mice. We measured locomotor activity in A53T mice using the Comprehensive Laboratory Animal Monitoring System (CLAMS). We found that A53T mice showed increased locomotor activity in the X, Y, and Z planes of the monitored cages (Figure 1D-F). Next, we checked the expression levels of αSyn in the whole brain of aged A53T mice. αSyn and the pathogenic phosphorylated pS129-αSyn were found to be upregulated in the brain of the A53T mice (Figure 1G-I). This was further confirmed by immunohistochemistry, which showed increased αSyn and pS129-positive staining in the cortex and striatum of A53T mice, indicating αSyn accumulation (Figure 1J-M). Tyrosine Hydroxylase (TH) is a rate-limiting enzyme responsible for converting L-tyrosine into L-DOPA for dopamine synthesis, and a marker for dopaminergic neurons. TH depletion is a hallmark of the nigrostriatal degeneration central to PD. We found decreased TH protein expression in the whole brain and decreased positive staining in the substantia nigra region in A53T mice compared to controls, suggesting dopaminergic neuron degeneration due to αSyn aggregation (Figure 1N-Q).

**Figure 1.**
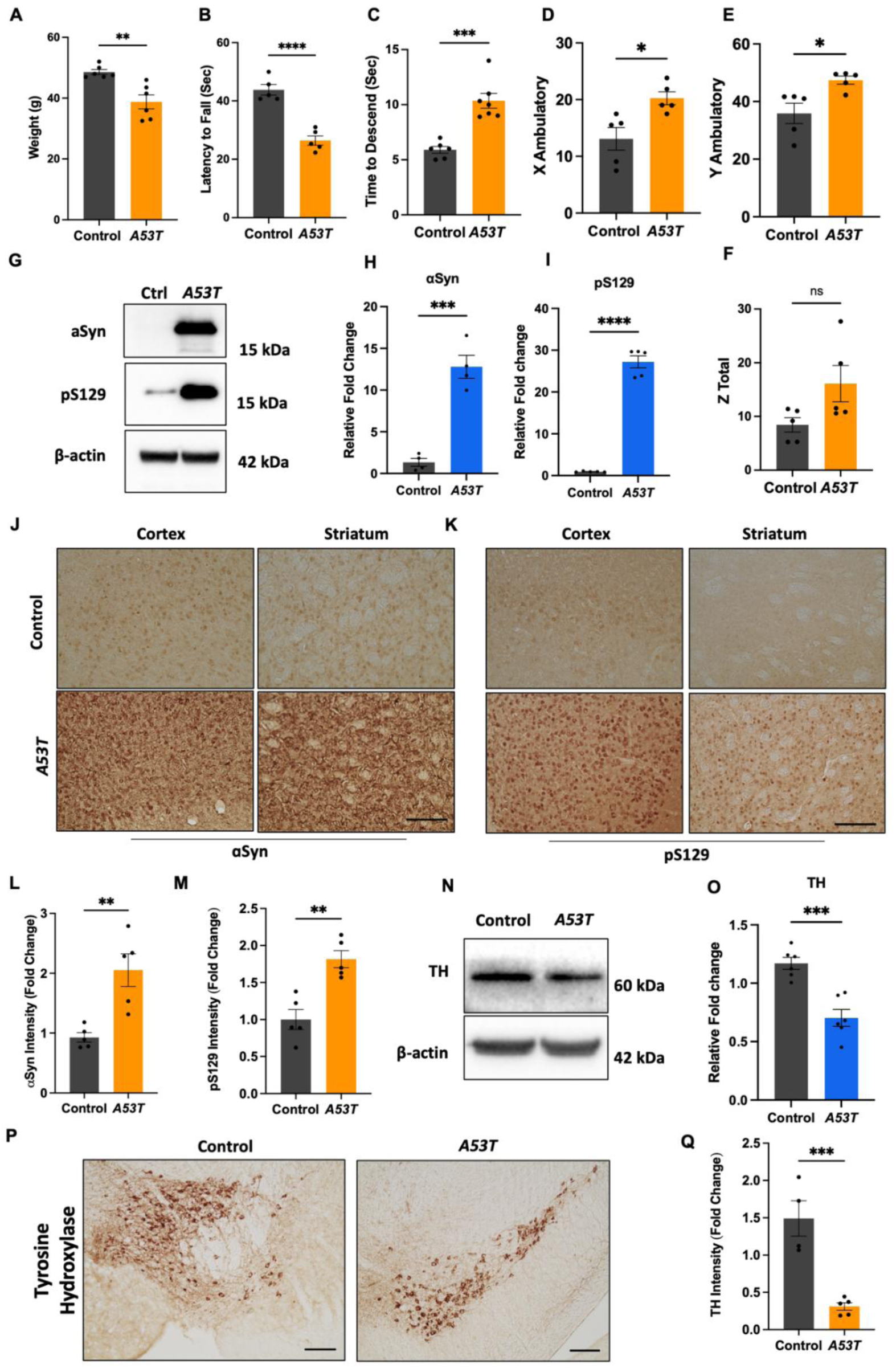
A53T mice exhibit motor coordination deficits, behavioral abnormalities, and neurodegeneration. A. Total body weight in grams of Control and human αSyn hemizygous (A53T^+/-^) mice measured at 18 months of age. B-F. Behavioral analysis of control and A53T mice at 18 months of age (B), Rotarod analysis (C), Pole Test (D-F), Total activity of mice measured in CLAMS. G-I. Representative western blot analysis for αSyn and pS129 and beta actin (loading control) using whole brain protein lysate (G) and their densitometric quantification (H, I). J-K. Representative micrographs of coronal brain sections stained for αSyn and pS129 showing cortex and striatum region (J, K) and their quantification as mean grey intensity (L, M) in control and A53T mice (n=6). N, O. Representative western blot analysis for Tyrosine Hydroxylase (TH) and beta actin (loading control) using whole brain protein lysate (N) and its densitometric quantification (O). P, Q. Representative micrographs of coronal brain sections stained for Tyrosine Hydroxylase at SNpc (P) and its quantification as mean grey intensity (Q) in control and A53T mice (n=6). Data presented as mean ± SEM; an unpaired, two-tailed Student’s t-test was employed, with \*\*\*\**P* < 0.0001, \*\*\**P* < 0.001, \*\**P* < 0.01, and \**P* < 0.05. Scale bar: 100 µm (J, K and P).

In the muscle, A53T mice exhibited reduced muscle weight specifically in the tibialis anterior (TA), gastrocnemius (GAS), quadriceps, and soleus muscles compared to age-matched control mice (Figure 2A-D). Because different muscles showed similar weight reduction trends, the gastrocnemius was used as a representative muscle for further study due to its relatively large size. The αSyn and pS129 protein levels were elevated in the gastrocnemius muscle (Figure 2E-G). These results indicate that the A53T mouse model of PD shows reduced muscle mass across different muscles, which may correlate with increased αSyn levels. To explore this, cross-sections of the gastrocnemius muscle were stained with laminin to visualise individual myofibers (Figure 2H). The myofibers were characterised by their cross-sectional area (CSA), grouped by size, and quantified (Figure 2I). We observed a decrease in the number of large myofibers and an increase in the proportion of small myofibers in the A53T mice. The total number of fibers per unit area was increased; however, the overall CSA and average fiber size decreased in the gastrocnemius muscle of A53T mice (Figure 2J-L). The A53T mice also exhibited reduced muscle function, as measured by the grip strength normalized to body weight (Figure 2M). This also correlates with the reduced motor coordination in the rotarod and pole test as discussed above. These results indicate that increased αSyn levels in the A53T PD mouse model are associated with neurological and muscular defects at the structural and functional levels.

**Figure 2.**
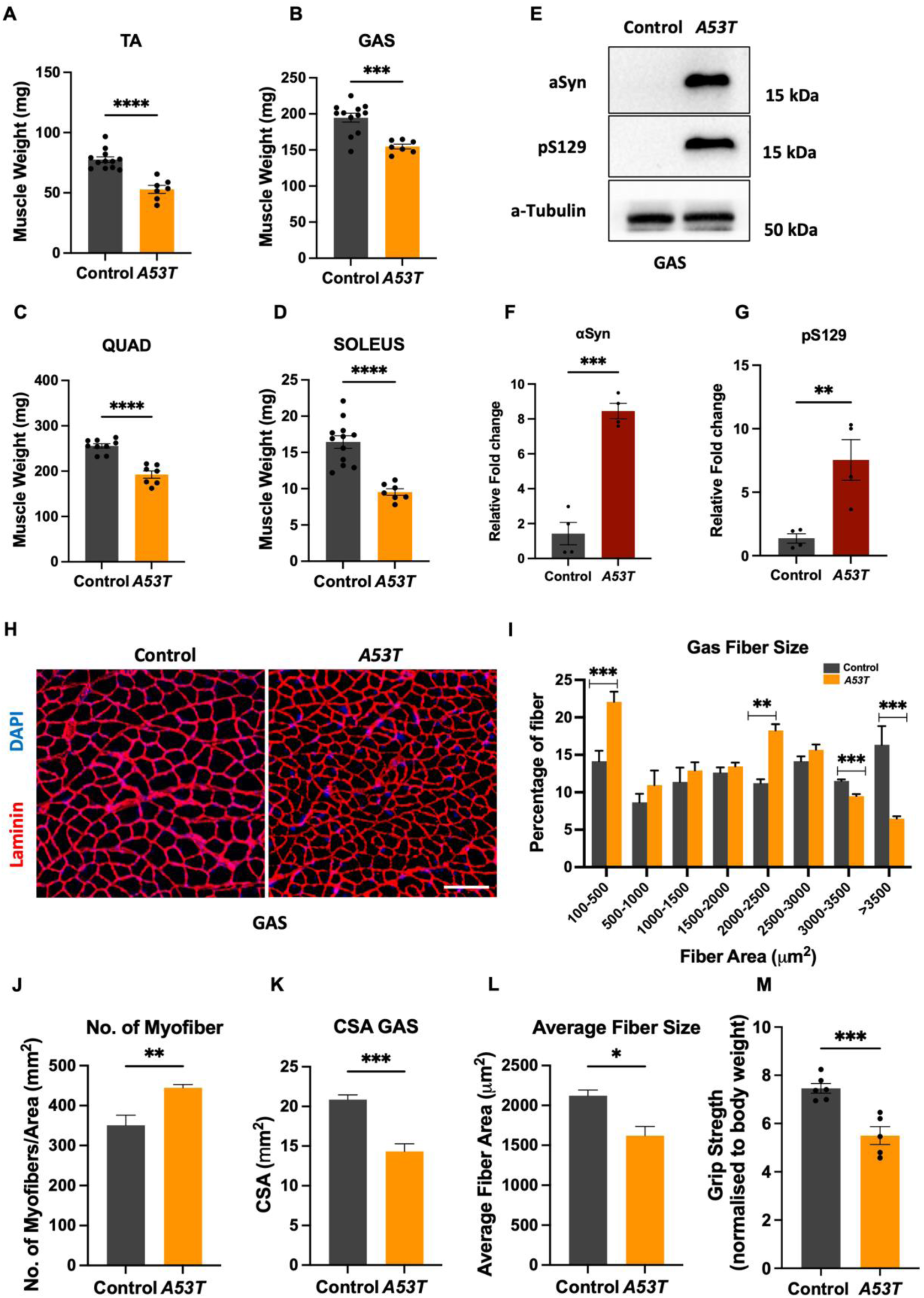
A53T mice exhibit loss of muscle mass along with structural and functional defects. A-D. Individual muscle weight (milligrams) of gastrocnemius (GAS), tibialis anterior (TA), quadriceps (Quad), and soleus muscle of control and A53T mice measured at 18 months of age. E-G. Representative western blot analysis for αSyn and pS129 and alpha tubulin (loading control) using protein lysates from the gastrocnemius muscle (E) and their densitometric quantification (F, G). H. Representative fluorescent micrographs of transverse sections through the gastrocnemius muscle of control and A53T mice stained with Laminin (red), and DAPI (blue). I. Quantification of the number of myofibers grouped according to myofiber area through the gastrocnemius muscle of control and A53T mice (n=6 mice per genotype). J-L. Quantification of the number of myofibers normalized to cross-sectional area (CSA) (mm2) (J), Total CSA (K), and average fiber size (L) through the gastrocnemius muscle of control and A53T mice. M. Quantification of grip strength control and A53T mice normalized to body weight (gf/gram body weight) at 18 months of age. Data presented as mean ± SEM; an unpaired two-tailed Student’s t-test was employed, with \*\*\*\**P* < 0.0001, \*\*\**P* < 0.001, \*\**P* < 0.01, and \**P* < 0.05. Scale bar: 100 µm (H).

### 2. A53T mouse model of PD exhibits neuromuscular and functional defects

We observed muscle defects, including myofiber changes and reduced grip strength in the A53T mice. Thus, we further investigated the NMJs in these mice. For this purpose, the gastrocnemius muscle was longitudinally sectioned and stained with α-bungarotoxin to mark the NMJs. α-bungarotoxin specifically labels acetylcholine receptors at the postsynaptic terminals of the NMJs. We found that αSyn was expressed in the NMJs of the A53T mice. Similarly, pS129 was detected specifically in the A53T mice compared to controls (Figure 3A). αSyn is a presynaptic protein and is reported to regulate synaptic vesicle trafficking and neurotransmitter release. As we found αSyn and pS129 enrichment in the NMJs, we were curious about NMJ-associated changes due to this enrichment. We stained the presynaptic and postsynaptic terminals of the NMJs with synaptophysin and α-bungarotoxin, respectively (Figure 3B). We observed no significant difference in the presynaptic or postsynaptic area occupied by the NMJs (Figure 3C, D). However, increased NMJ fragmentation was seen in the gastrocnemius muscle in the A53T mice (Figure 3E). We also observed an increase in the number of nuclei per NMJ, suggesting a compensatory pathological response (Figure 3F) [44]. To investigate NMJ molecular changes, we assessed key synaptic proteins in the gastrocnemius muscle of 18-month-old control and A53T mice (Figure 3G-K). We found that NCAM1, a denervation marker [45,46], was significantly downregulated in A53T mice, suggesting failed regeneration due to chronic αSyn toxicity overwhelming motor neurons[47,48]. The acetylcholine receptor (AChR) protein levels were increased, indicating compensation for reduced NCAM1 function in the A53T mice. Postsynaptic agrin signaling regulates presynaptic AChR clustering. Agrin from motor neurons binds LRP4, activating MuSK to cluster AChRs, thus maintaining synaptic stability[49,50]. We found reduced LRP4 and MuSK protein levels, suggesting impaired agrin signaling, explaining the compensatory increase in AChR levels. We also performed an ex vivo muscle contractility assay on the gastrocnemius muscle from the A53T and control mice (Figure 3L, M). Consistent with the molecular perturbations, upon electrical stimulation at increasing frequencies, the A53T muscle of A53T mice showed markedly reduced contractile force and maximum contractile force generation compared with controls, suggesting impaired synaptic transmission and neuromuscular efficacy. These findings demonstrate that pathological αSyn accumulation at the NMJs leads to NMJ fragmentation, altered agrin signaling, and ex vivo contractile deficits, indicating ineffective synapses in the A53T mice.

**Figure 3.**
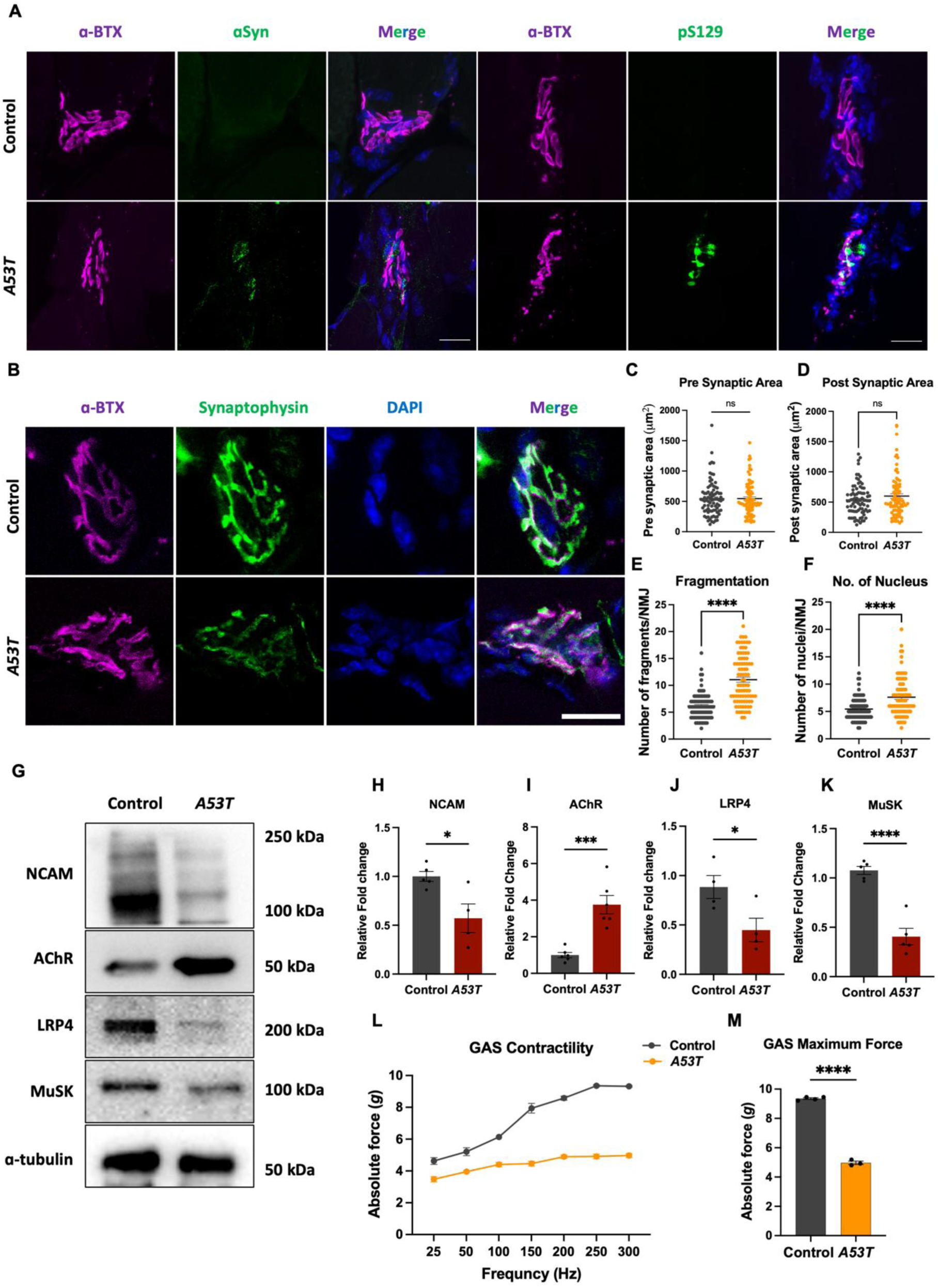
A53T mice exhibit NMJ fragmentation, degeneration, and contractility defects. A. Representative fluorescent micrographs of longitudinal sections through the gastrocnemius muscle of 18-month-old control and A53T mice labeled for α-Bungarotoxin (pink), αSyn and pS129 (green), and DAPI (blue). B-F. Representative fluorescent micrographs of longitudinal sections through the gastrocnemius muscle of 18-month-old control and A53T mice labeled for α-Bungarotoxin (pink), synaptophysin (green), and DAPI (blue) (B). Quantification of the number of pre-synaptic area (C), Post-synaptic area (D), fragments/neuromuscular junction (NMJ) (E), and Number of nuclei/NMJ (F). G-K. Representative western blot analysis for NCAM, AChR, LRP4, MuSK, and alpha tubulin (loading control) using protein lysates from the gastrocnemius muscle (G) and their densitometric quantification (H, K). L-M. Quantification of muscle contractility (L) and maximum force generation (M) in the gastrocnemius muscle of control and A53T mice at 18 months of age (n=3 mice per genotype). Data presented as mean ± SEM; an unpaired two-tailed Student’s t-test was employed, with \*\*\*\**P* < 0.0001, \*\*\**P* < 0.001, \*\**P* < 0.01, and \**P* < 0.05. Scale bar: 100 µm (A), 10 µm (B).

### 3. A53T mice model shows systemic inflammation signatures in the plasma

PD is increasingly recognized as a multisystem disorder in which pathology extends beyond the central nervous system to peripheral tissues. A major contributor to this systemic involvement is immune activation, where neurodegeneration triggers microglial responses that drive neuroinflammation and promote the release of circulating cytokines. These inflammatory mediators can propagate to peripheral tissues and, in turn, further exacerbate neurodegenerative processes. To determine whether this systemic inflammatory signature is recapitulated in the A53T mouse model, we profiled plasma proteins using targeted Multiple Reaction Monitoring (MRM) mass spectrometry. MRM provides sensitive, multiplexed detection and relative quantitation of low-abundance proteins, including cytokines, chemokines, and acute-phase proteins, in circulation, enabling correlation of both central neurodegeneration and peripheral muscle alterations. We designed an MRM workflow for several proteins selected from the PD literature and analysed their differential expression pattern using a targeted pipeline (Figure S1A). MRM analysis revealed a distinct pattern of innate immune activation in the A53T mice compared to controls (Figure S1B, C). We found significant upregulation of pro-inflammatory cytokines, namely Tumor Necrosis Factor (TNF) and Interleukin-1 beta (IL-1β), reflecting broad innate inflammatory cytokine signaling. Similarly, Interleukin-18 (IL-18) upregulation suggested inflammasome activation as a contributing mechanism. Complement pathway engagement was evidenced by increased circulating levels of Complement component 3 (C3), suggesting complement activation. Notably, GFAP (Glial Fibrillary Acidic Protein), a structural protein canonically associated with astrocytes[51], was also significantly elevated in the plasma, suggesting active CNS inflammation and blood-brain barrier compromise, leading to the release of glial proteins into circulation. Elevated CX3CL1 (fractalkine), a chemokine with roles in leukocyte recruitment and microglia-neuron communication[52], further indicated active neuroimmune crosstalk extending to the periphery. Similarly, circulating C-reactive protein (CRP), a classical hepatic acute-phase reactant[53], was significantly elevated, indicating a robust liver-driven acute-phase response secondary to innate immune activation. Taken together, A53T mice exhibit robust systemic inflammation, with bidirectional signaling between the central nervous system and the peripheral immune compartment, further exacerbating neurodegeneration. Overall, the PD mouse model exhibits hallmark motor deficits accompanied by brain pathology, muscle atrophy, and systemic inflammation.

### 4. A53T PD mice exhibit proteome-wide alterations in the brain and muscle

The parallel brain pathology and muscle atrophy, along with neuromuscular defects and robust systemic inflammation observed in the A53T mice, suggest potential brain-muscle crosstalk contributing to PD progression, though the underlying molecular mechanisms remain undefined. Thus, we aimed to identify shared molecular pathways or proteins underlying PD progression in these two tissues. To address this, we performed label-free SWATH-MS proteomics on the brain and gastrocnemius muscle tissues from 18-month-old A53T mice and littermates. Mass spectrometry data acquisition achieved an overall data completeness of ∼ 98.2 % in the brain samples and ∼94.3 % in the muscle at the protein group level. The average peptide quantified per protein group was 7.6 and 8.2 for brain and muscle, respectively. The median protein coefficient of variation (%CV) in the control and A53T mice brain data was 10.7 % and 10.8 %, respectively, whereas, in the muscle samples, the median %CV was 19.3 % and 14.8 % in the control and A53T groups, respectively. We identified 3090 proteins in the brain and 1853 in the muscle, of which 1370 proteins were found to be common between the brain and muscle proteome (Figure 4A) (Table S2, S3). Principal component analysis (PCA) separated the control and A53T mice samples into two distinct but partially overlapping islands, with a maximum variation of 45.65 % at PC1 and 14.26 % at PC2 in the brain tissue (Figure 4B). In muscle samples, a similar pattern of sample clustering was observed in PCA, with maximum variation of 45.65 % at PC1 and 14.92 % at PC2 (Figure 4C). The identified proteins were visualised using a heatmap, which highlighted regions of differential protein expression between controls and the A53T mice (Figure 4D, E). The regulated brain and muscle proteins between the control and the A53T mice were filtered based on an absolute log2 FC cutoff of 0.58 and FDR-adjusted p<0.05. We mapped 27 upregulated and 41 downregulated proteins in the brain proteome (Figure 4F). However, in the muscle tissue, 113 proteins were upregulated, and 48 were downregulated in the A53T mice (Figure 4G). Among the upregulated proteins, 4 were common to both brain and muscle, 23 were unique to the brain, and 107 were unique to muscle. No common proteins were down-regulated among the two tissues (Figure 4H). In both brain and muscle tissues, αSyn was uniquely upregulated in the A53T mice, confirming the overexpression of αSyn in both the brain and the skeletal muscle. Next, we performed pathway analysis to identify the functional role of the dysregulated proteins in the brain and the muscle of the A53T mice compared to controls. For this purpose, regulated proteins were used for pathway analysis against the Gene Ontology Biological Process (GOBP) database, and the top-ranked pathways were selected (Figure S2A, B). In the brain, we found pathways related to neurodegeneration, neuroactive ligand receptor interactions, neurotransmitter release, and synaptic vesicle cycles were uniquely enriched. In the gastrocnemius muscle, the pathways related to aerobic respiration, HIF-1 signalling, and mitochondrion organization were enriched. Interestingly, we also found neuromuscular pathways to be dysregulated in the A53T mice, further strengthening our above finding of neuromuscular defects in these mice. Next, we focused on the common pathways dysregulated in both the brain and the muscle tissues of the A53T mice. We found dysregulated pathways related to Parkinson’s disease, oxidative stress, ion transport, oxidative phosphorylation, cell death, and cellular stress response were shared between the brain and the muscle. These results suggest that αSyn-induced molecular alterations lead to PD development in the A53T mice. Among the common pathways identified, the iron uptake and transport and ferroptosis pathways were particularly enriched in the brain and muscle, respectively. We became interested in further exploring iron-related pathways, as iron dysregulation driving neuronal ferroptosis is well documented in the PD brain; however, iron dysregulation and skeletal muscle ferroptosis have not been reported in the PD skeletal muscle.

**Figure 4.**
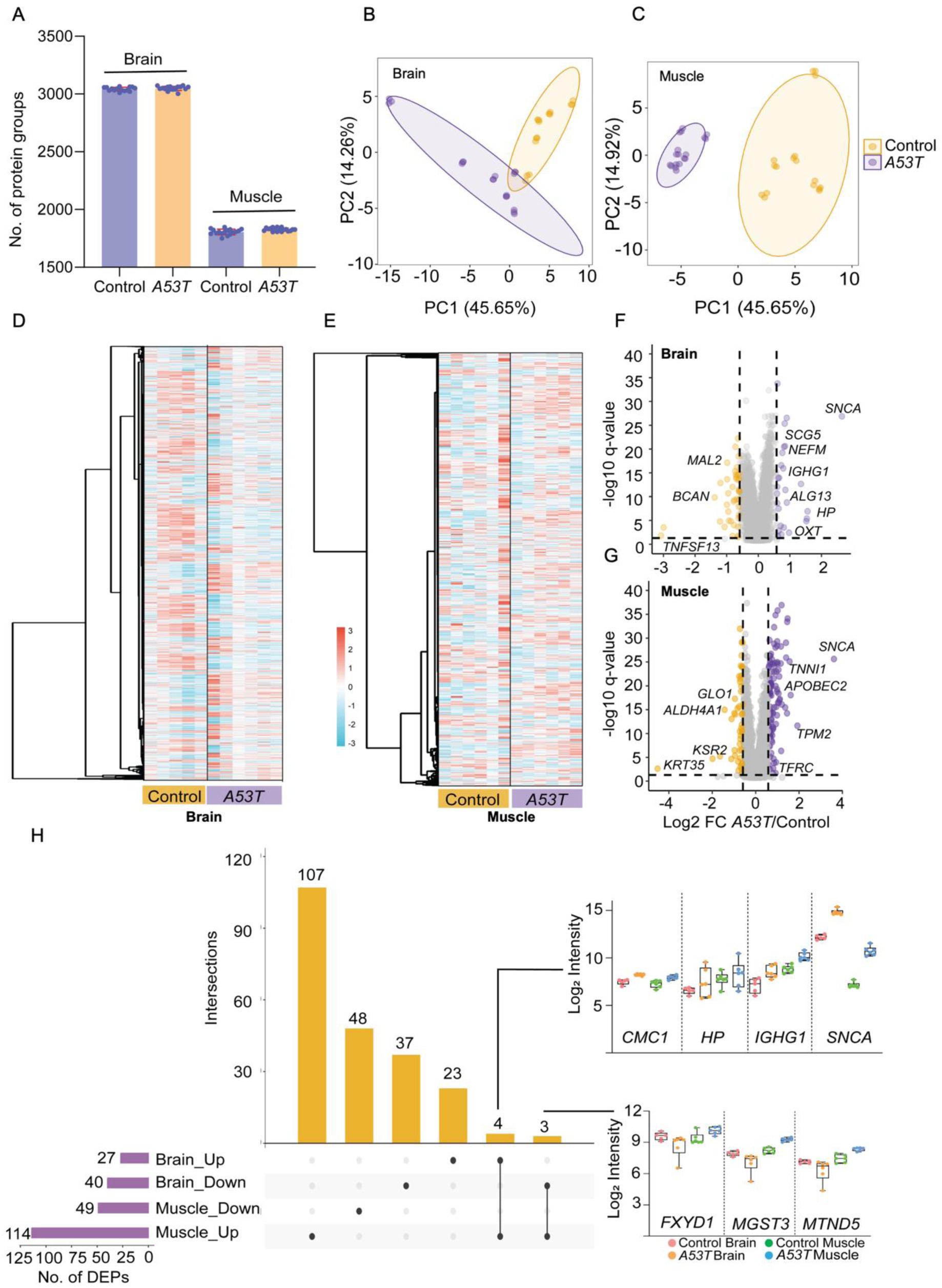
Proteome profiling of the brain and skeletal muscle of A53T and control mice. A. The representation of the number of protein groups identified in the brain and gastrocnemius muscle of control and A53T mice (Brain, control n=5, A53T n=6) and (Muscle, control and A53T n=6). B, C. Principal Component Analysis (PCA) of the brain and muscle samples from control and A53T mice. The PC1 and PC2 variance were plotted on x and y-axis respectively. D, E. Heat maps show the distribution of identified proteins in the brain (D) and gastrocnemius muscle (E) across samples. F, G. The volcano plot highlights regulated proteins between A53T and control, with log2 fold change (FC) and -log10 q-value plotted on x and y-axis, respectively, in the brain (F) and gastrocnemius muscle (G). The purple and yellow circles denote the respective up- and downregulated proteins, respectively. The horizontal dotted line represents the -log10 (q) value cutoff of 1.3, whereas vertical lines mark the absolute log_2_ FC cutoff of 0.58 in opposite directions. H. The upset plot shows the overlap between regulated proteins identified in the control vs A53T comparison in brain and gastrocnemius muscle samples, with the intensity distribution of common proteins shown in the right panel. Statistical evaluation of the DEPs was done using an unpaired two-tailed Student’s t-test with multiple testing correction.

We further explored the key proteins indicating iron dysregulation and ferroptosis in our PD proteomics data. We searched the identified proteins against the ferroptosis database, FerrDb V3, to identify ferroptosis markers in the brain and muscle proteomes [54], where 46 and 36 ferroptosis-associated proteins were identified in the brain and muscle, respectively. 29 proteins were common to both organs, whereas 17 and 7 proteins were unique to the brain and muscle, respectively (Figure S3A) (Table S4, S5). The regulated brain and muscle ferroptosis proteins between the control and the A53T mice were narrowed down based on an absolute log2 FC and significance (Figure 5A, B). The identified proteins were visualised using a heatmap, which highlighted regions of differential protein expression between control and A53T mice (Figure 5C, D). We further segregated the identified proteins into ferroptosis markers, suppressors, and drivers using the FerrDb V3 database (Figure S3B, C). In total, 19 ferroptosis suppressors were detected in both the brain and the skeletal muscle, including ALDH2, FTL1, FXN, OTUB1, PARK7, PRDX6, CISD2, DHODH, ENO1, G3BP1, GOT1, HSPA8, HSPB1, IDH2, NFS1, PCBP1, TIGAR, TMSB4X, and TXNDC12. These proteins form a shared anti-ferroptotic network that induces iron sequestration (FTL1, PCBP1, FXN, NFS1), maintains mitochondrial redox homeostasis (IDH2, DHODH, ALDH2), supports lipid peroxide detoxification (PRDX6), and modulates cellular stress responses (PARK7, HSPA8, HSPB1).

**Figure 5.**
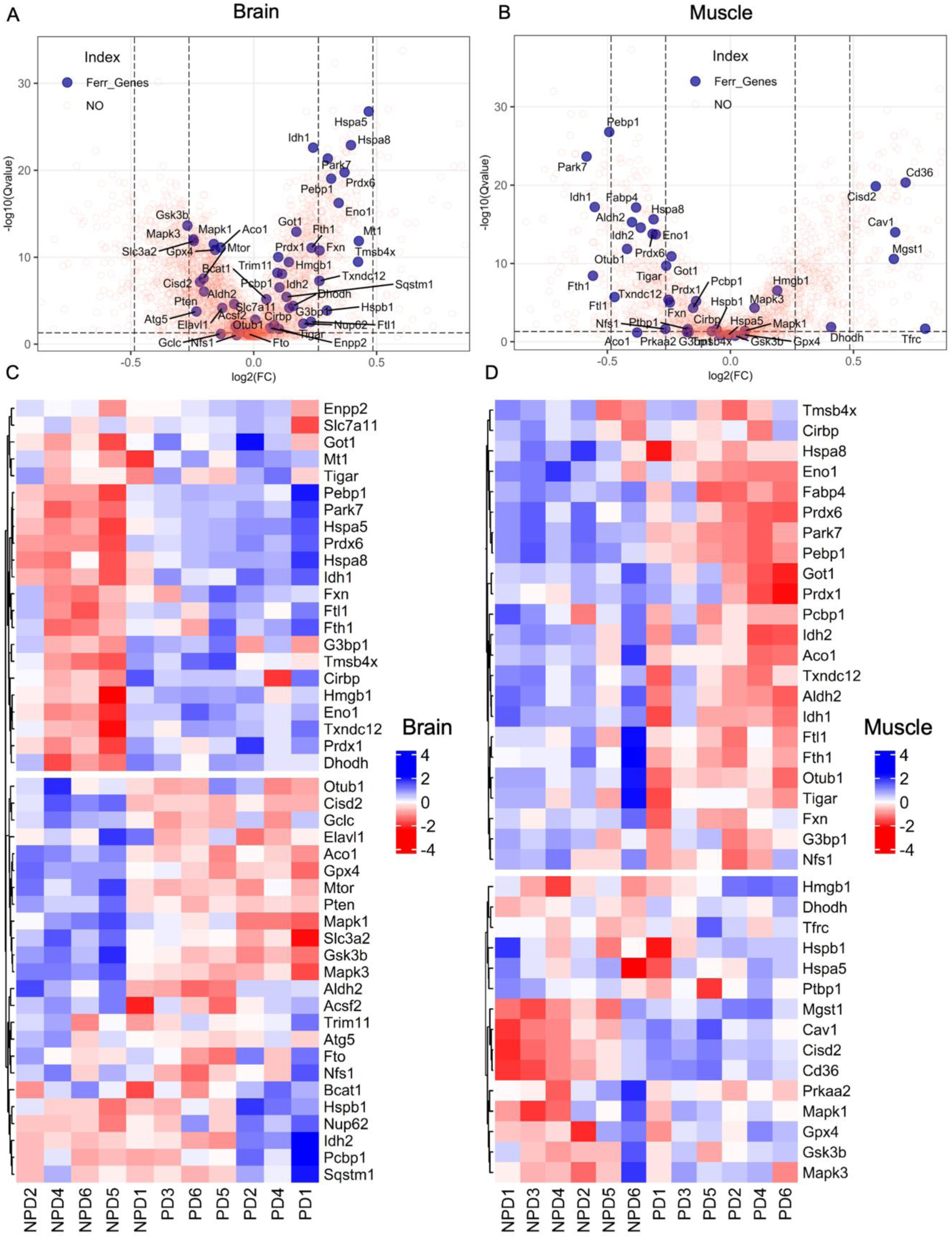
Ferroptosis-associated proteins were enriched in the brain and the skeletal muscle of the A53T mice model of PD. A, B. The volcano plot comparing ferroptosis-associated proteins between A53T and control, with log2 fold change (FC) on the x-axis and -log10 q-value on the y-axis, in the brain (A) and in the gastrocnemius muscle (B). The purple color depicts ferroptosis-associated proteins, while the black dotted lines show the FC and significance cut-off. C, D. Z-score abundance distribution of ferroptosis-associated proteins were plotted as heat map, which were identified in the brain (C) and gastrocnemius muscle (D) samples across the comparison groups. Statistical evaluation of the DEPs was done using an unpaired two-tailed Student’s t-test with multiple testing correction.

In addition, six ferroptosis driver proteins, namely ACO1 (IRP1), GSK3B, CIRBP, HMGB1, IDH1, and MAPK3, were common to both tissues, indicating ongoing stress and active iron-responsive signaling pathways that may induce lipid peroxidation and ferroptotic sensitivity. Consistent with the identification of canonical ferroptosis pathway proteins, two hallmark ferroptosis markers, GPX4 and FTH1, were detected in both organs, whereas SLC7A11 and TFRC were unique to the brain and the skeletal muscle, respectively. Thus, these markers reflect an active role of iron uptake, storage, and phospholipid peroxide detoxification in regulating ferroptosis in the two organs. Together, these findings indicate that the brain and the skeletal muscle of the A53T PD mouse model exhibit dysregulated iron homeostasis and ferroptosis.

### 5. Ferroptosis is a shared molecular mechanism that drives PD pathology in the brain and muscle of the A53T mouse model

To validate our mass spectrometry data showing alterations in iron homeostasis in PD mice, we assessed the expression levels of proteins involved in iron regulation. For this purpose, brain and muscle protein lysate from 18-month-old A53T and control mice was used, followed by western blotting and densitometry analysis. We assessed the levels of key proteins involved in the iron regulation pathway, including the iron uptake receptor Transferrin receptor (TFRC), the endosomal iron transporter Divalent metal ion transporter (DMT1), and the iron storage protein Ferritin Heavy Chain 1 (FTH1) (Figure 6A-C). In the brain, TFRC, DMT1, and FTH1 protein levels were significantly increased in the A53T mice compared with controls, indicating coordinated activation of iron uptake and sequestration pathways. However, in the muscle, although TFRC protein expression was increased, we did not observe significant changes in DMT1 levels. Notably, FTH1 protein levels were significantly reduced in the skeletal muscle, indicating impaired iron storage capacity despite increased iron influx. Reduced ferritin levels may reflect increased ferritin turnover or degradation, potentially mediated through ferritinophagy pathways involving NCOA4, thus promoting labile iron accumulation[55]. To further investigate whether these molecular changes lead to alterations in total iron content, we measured total iron levels in the brain and skeletal muscle homogenates (Figure 6D, E). We found a significant increase in total iron levels in the brains and skeletal muscles of A53T mice, demonstrating systemic iron accumulation in the PD model. Together, these findings suggest that increased TFRC-mediated iron uptake contributes to iron overload in both the brain and muscle tissues, while the downstream iron sequestration process shows tissue-specific alterations.

**Figure 6.**
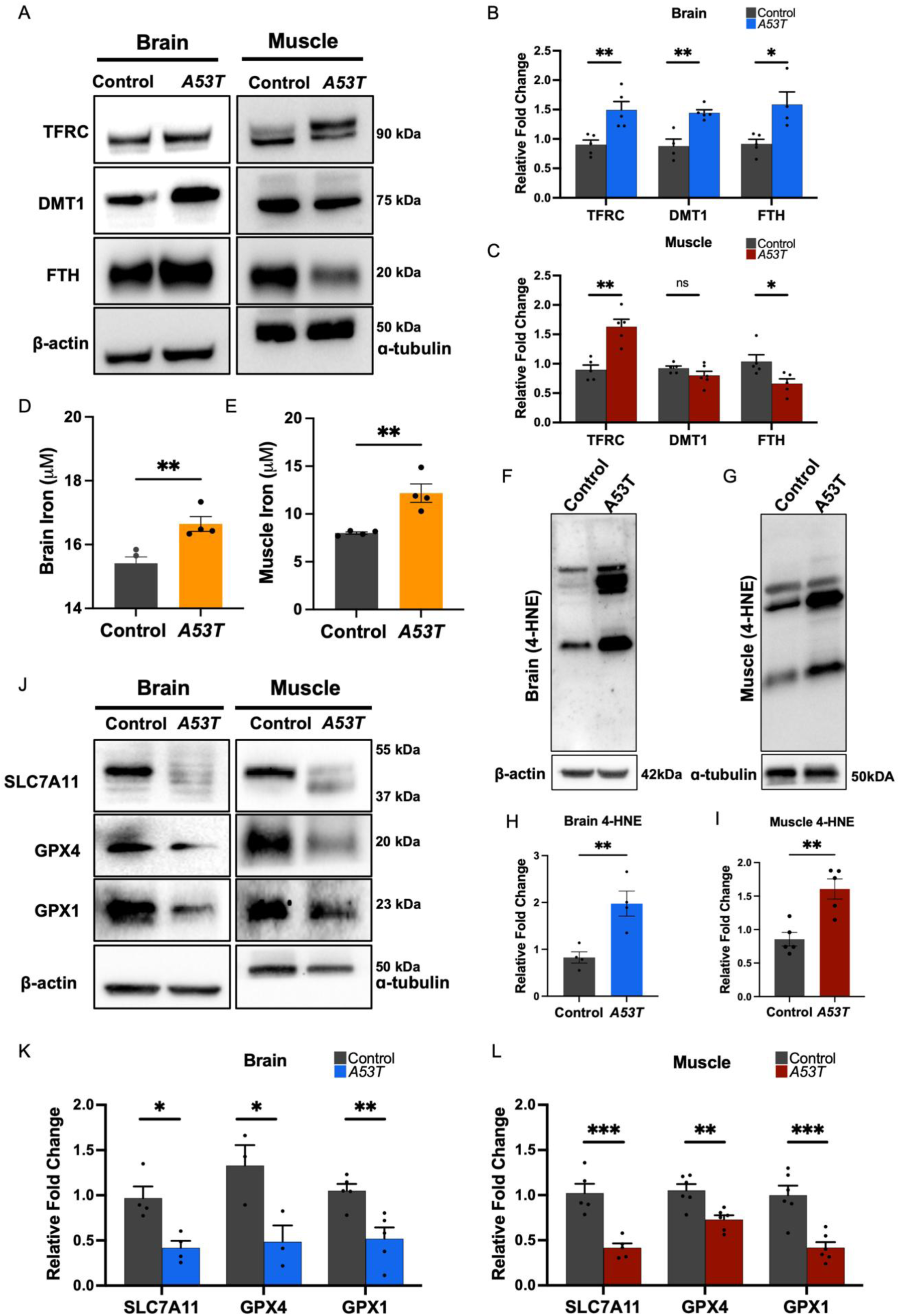
Dysregulated iron homeostasis in the brain and skeletal muscle of A53T mice. A-C. Representative western blot analysis for TFRC, DMT1, FTH, and beta-actin and alpha tubulin (loading control) using protein lysates from the brain and gastrocnemius muscle (A) and their densitometric quantification (B, C). D, E. Total iron measurement from the tissue lysate of brain (D) and gastrocnemius muscle (E) from A53T and control mice. F-I. Representative western blot analysis for 4-HNE, beta-actin, and alpha tubulin (loading control) using protein lysates from the brain (F) and gastrocnemius muscle (G), and their densitometric quantification (H, I), respectively. J-L. Representative western blot analysis for SLC7A11, GPX4, GPX1, and beta-actin, and alpha tubulin (loading control) using protein lysates from the brain and gastrocnemius muscle (J), and their densitometric quantification (K, L), respectively. Data presented as mean ± SEM; an unpaired two-tailed Student’s t-test was employed, with, \*\*\**P* < 0.001, \*\**P* < 0.01, and \**P* < 0.05.

Increased intracellular iron is known to promote oxidative damage through iron-dependent lipid peroxidation, a key driver of ferroptosis[31]. To assess lipid peroxidation, we measured levels of 4-hydroxynonenal (4-HNE) in the brain and skeletal muscles of the A53T mice (Figure 6F-I). 4-HNE is a key byproduct of lipid peroxidation, primarily from the oxidative breakdown of ω-6 polyunsaturated fatty acids (PUFAs) such as linoleic and arachidonic acids in cell membranes[56]. Western blot analysis revealed significantly elevated 4-HNE levels in both the brain and skeletal muscle of A53T mice, indicating enhanced lipid peroxidation in both tissues. Under physiological conditions, lipid peroxides are reduced by glutathione-dependent antioxidant defenses, thereby limiting ferroptotic damage[57–60]. Consistent with our proteomic data indicating dysregulation of glutathione metabolism, we next examined the expression of key antioxidant regulators, including the cystine/glutamate antiporter SLC7A11 (xCT) and the glutathione peroxidases Glutathione peroxidase 4 (GPX4) and Glutathione peroxidase 1 (GPX1) (Figure 6J-L). Western blot analysis demonstrated that SLC7A11, GPX4, and GPX1 protein levels were significantly decreased in both the brain and skeletal muscle of A53T mice. Collectively, these findings indicate that iron accumulation in both brain and skeletal muscle is accompanied by increased lipid peroxidation and impaired antioxidant defenses, increasing ferroptotic susceptibility. These results support the presence of shared ferroptosis-associated mechanisms across the neuromuscular axis in the A53T PD model, potentially linking central neurodegeneration with peripheral muscle dysfunction.

### 6. NRF2 signaling and antioxidant defenses are suppressed in the brain and skeletal muscle of A53T mice

We observed increased iron accumulation, lipid peroxidation, and suppression of glutathione-dependent antioxidant defenses, and sought to determine whether broader oxidative stress response pathways were compromised in the A53T mice. Our proteomic analysis also revealed significant enrichment of dysregulated oxidative stress and ROS pathways, prompting us to assess the Nuclear factor erythroid 2–related factor 2 (NRF2) signaling pathway. NRF2 is a central transcriptional regulator of cellular antioxidant defenses and a known modulator of ferroptosis susceptibility [30,61,62]. We found a significant reduction in total NRF2 and phosphorylated NRF2 (p-NRF2) protein levels by western blot analysis in both the brain (Figure S4A, C) and skeletal muscle (Figure S4B, D) of A53T mice. The NRF2 levels are regulated by KEAP1 and Sequestosome 1 (SQSTM1/p62) proteins. KEAP1 interacts with NRF2 and targets it for degradation via the ubiquitination-proteasome pathway. p62/SQSTM1 stabilizes NRF2 by disrupting its interaction with KEAP1, thereby promoting NRF2 nuclear accumulation[63–65]. In both the brain (Figure S4A, C) and muscle (Figure S4B, D) of the A53T mice, we observed a significant decrease in p62 expression; however, KEAP1 levels remained unchanged. The reduction in p62 levels, therefore, suggests impaired NRF2 activation and its downstream transcriptional target antioxidant genes. This finding is in agreement with our observation that SLC7A11 and GPX1, the targets of NRF2, are downregulated in the brain and skeletal muscle of A53T mice. We also checked the status of the antioxidant enzymes, namely, Catalase, Superoxide dismutase 1 (SOD1), Peroxiredoxin 2 (PRDX2), and Peroxiredoxin 4 (PRDX4), playing essential roles in detoxifying reactive oxygen species and protecting cells from oxidative damage[66–69]. In addition to suppression of NRF2 signaling, we found significant downregulation of Catalase, SOD1, PRDX2, and PRDX4 protein levels in both the brain (Figure S4E, G) and the skeletal muscle of the A53T mice (Figure S4F, H). Their coordinated reduction, therefore, indicates a global impairment of antioxidant capacity in both the brain and skeletal muscle. Taken together with our earlier findings of iron overload and increased lipid peroxidation, these results further suggest impaired antioxidant and ferroptosis-protective responses in both the brain and the skeletal muscle of the A53T mice.

### 7. αSyn interacts with TFRC and remodels the cell-surface proteome to promote ferroptosis-associated signaling

As we discussed, iron overload, lipid peroxidation, and impaired NRF2-dependent antioxidant defenses contribute to increased ferroptosis susceptibility in the brain and skeletal muscle of the A53T mouse model of PD. We next sought to identify upstream mechanisms linking αSyn pathology to ferroptotic signaling. We investigated whether extracellular αSyn aggregates alter the surface proteome of neuronal and muscle cells. Because our PD model expresses mutant human A53T αSyn, we hypothesized that αSyn aggregates may interact with cell-surface receptors to initiate signaling pathways that regulate iron metabolism and ferroptosis.

To investigate this, we performed cell-surface proteomics in mouse neuroblastoma (N2A) and mouse myoblast (C2C12) cells following exposure to αSyn aggregates (Figure 7A). For this purpose, αSyn protein was purified, and preformed fibrils (PFFs) were prepared as previously described [20]. The cells were treated with αSyn-PFFs or PBS (vehicle control) and incubated for 2h. Cell-surface proteins were selectively labeled with membrane-impermeable NHS-LC biotin. Biotinylated proteins were captured by streptavidin pull-down, followed by mass spectrometry analysis. The identified proteins were filtered for biotin modifications and surface localization using the Cell Surface Protein Atlas (CSPA) [70]. Upon exposure to αSyn, we found several cell-surface receptors to be significantly altered. In N2A cells, we identified 36 surface proteins, of which 24 were common to both αSyn treatment and control (Figure 7B) (Table S6), whereas 9 and 3 were unique to αSyn treatment and control, respectively. Similarly, in C2C12 cells, we identified 55 surface proteins, of which 37 were common to both αSyn treatment and control (Figure 7C) (Table S7), whereas 1 protein was unique to αSyn treatment and 17 were unique to control, respectively. Next, we searched for proteins that were differentially regulated in αSyn PFF-treated conditions compared to controls (Figure 7D, E). In N2A cells, we identified 2 proteins, TFRC and VDAC1, to be upregulated, whereas RPN2 was downregulated. In C2C12 cells, we identified 9 proteins, namely, TFRC, NCAM1, STIM1, PLPP1, LAMP1, CKAP4, ENO1, HNRNPK, and SIDT2, as significantly upregulated and 2 proteins, FBN1 and ITGAV, as downregulated, respectively. The surface proteome data identified several novel cell-surface receptors that have not been explored previously in the context of αSyn. As we focused on identifying surface proteins that mediate ferroptosis, we identified a few candidate proteins with known or emerging roles in iron metabolism and ferroptosis regulation, notably, TFRC, responsible for transferrin-mediated iron uptake [71,72], stromal interaction molecule 1 (STIM1), a Ca^2+^ sensor, which has been recently shown to interact with TFRC and promote ferroptosis[73], voltage-dependent anion channel 1 (VDAC1) shown to induce ferroptosis by modulating mitochondrial ferritin[74] and α-Enolase 1 (ENO1) that suppresses cancer cell ferroptosis by degrading the mRNA of iron regulatory protein 1[75], among others. Importantly, our surface proteomics data identify TFRC as a common cell-surface protein upregulated in both N2A and C2C12 cells upon αSyn-PFF treatment (Figure 7F). These results are consistent with our in vivo SWATH-MS proteomic data, which showed increased TFRC expression in the skeletal muscle of 18-month-old A53T PD mice, as well as our biochemical findings demonstrating elevated TFRC expression and iron accumulation in brain and muscle tissues. To investigate whether αSyn directly interacts with TFRC, we fluorescently labelled αSyn with NHS-Rhodamine and prepared αSyn-PFFs. We treated N2A and C2C12 cells with labeled αSyn-PFFs and performed confocal microscopy, and found that αSyn colocalizes with TFRC in both N2A and C2C12 cells (Figure 8A, B). We further sought to understand the specificity of this colocalization; hence, we blocked surface TFRC with an anti-TFRC antibody, where anti-IgG served as an isotype control. We treated the N2A and C2C12 cells with labeled αSyn-PFFs, and confocal microscopy was performed to measure the fluorescence intensity of labelled αSyn-PFFs at the cell surface (Figure 8C, D). We found that surface blocking of TFRC with an anti-TFRC antibody significantly reduced αSyn-PFFs at the cell surface compared to treatment with αSyn-PFF alone. We did not observe a significant reduction in αSyn-PFFs fluorescence intensity in anti-IgG-treated cells (Figure 8E, F). These results indicate the specificity of αSyn towards TFRC. To further characterize this interaction, we performed biolayer interferometry (BLI) to quantify the interaction between αSyn and TFRC. For this purpose, recombinant human TFRC was immobilized to AR2G sensors, and purified human αSyn protein was kept in solution. The association and dissociation reactions were performed to check the interaction and to determine binding kinetics (Figure 8G, H). We found that αSyn binds hTFRC with a dissociation constant *K*_d_ of 360 nM. As expected, recombinant transferrin bound TFRC with a significantly higher affinity, *K*_d_ **=** 3.4 nM, serving as a positive control for TFRC ligand binding. Together, these findings indicate that TFRC is an αSyn-interacting receptor and suggest a potential mechanism by which extracellular αSyn aggregates promote iron uptake and increase ferroptosis susceptibility.

**Figure 7.**
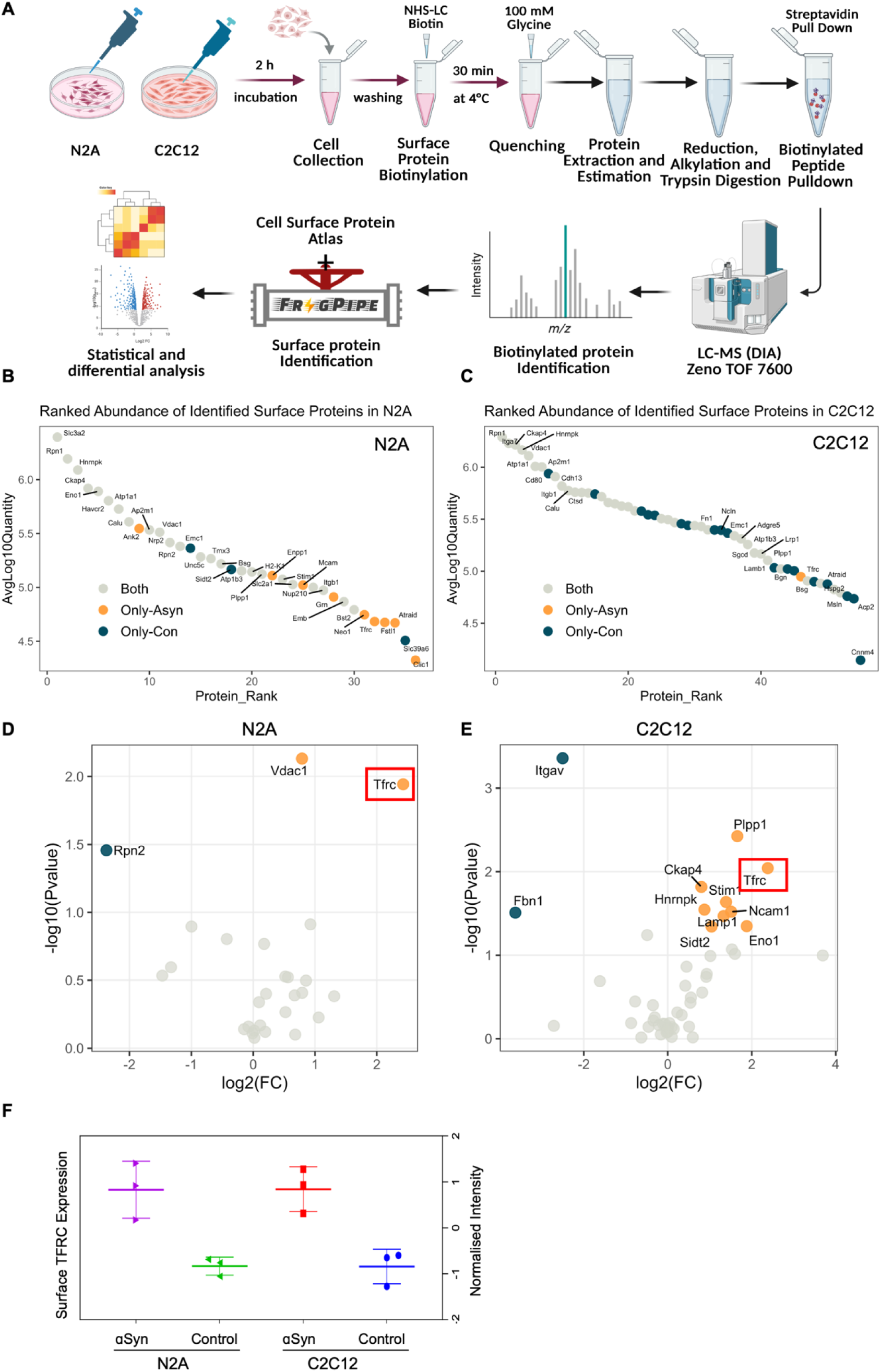
α-Synuclein remodels the surface proteome of brain and muscle cells and targets TFRC. A. Schematic workflow of cell surface protein labeling upon αSyn PFF treatment in N2A and C2C12 cells, followed by LC-MS analysis (n=3). B, C. Rank abundance plot for surface proteins identified in N2A (B) and C2C12 (C) cells. The protein abundance rank on the x-axis is plotted against its average log10 quantity on the y-axis, and the dots are colored according to identification in a comparison groups. D, E. Volcano plot comparing surface proteins between αSyn-PFF and control treatment, with log2 fold change (FC) on the x-axis and -log10 q-value on the y-axis, in N2A (D) and in the C2C12 cells (E). The up and downregulated proteins are labelled and denoted with orange and dark-green colour respectively. F. Box plot depicting surface TFRC expression in N2A and C2C12 cells upon αSyn-PFF and control treatment, plotted against normalised intensity.

**Figure 8.**
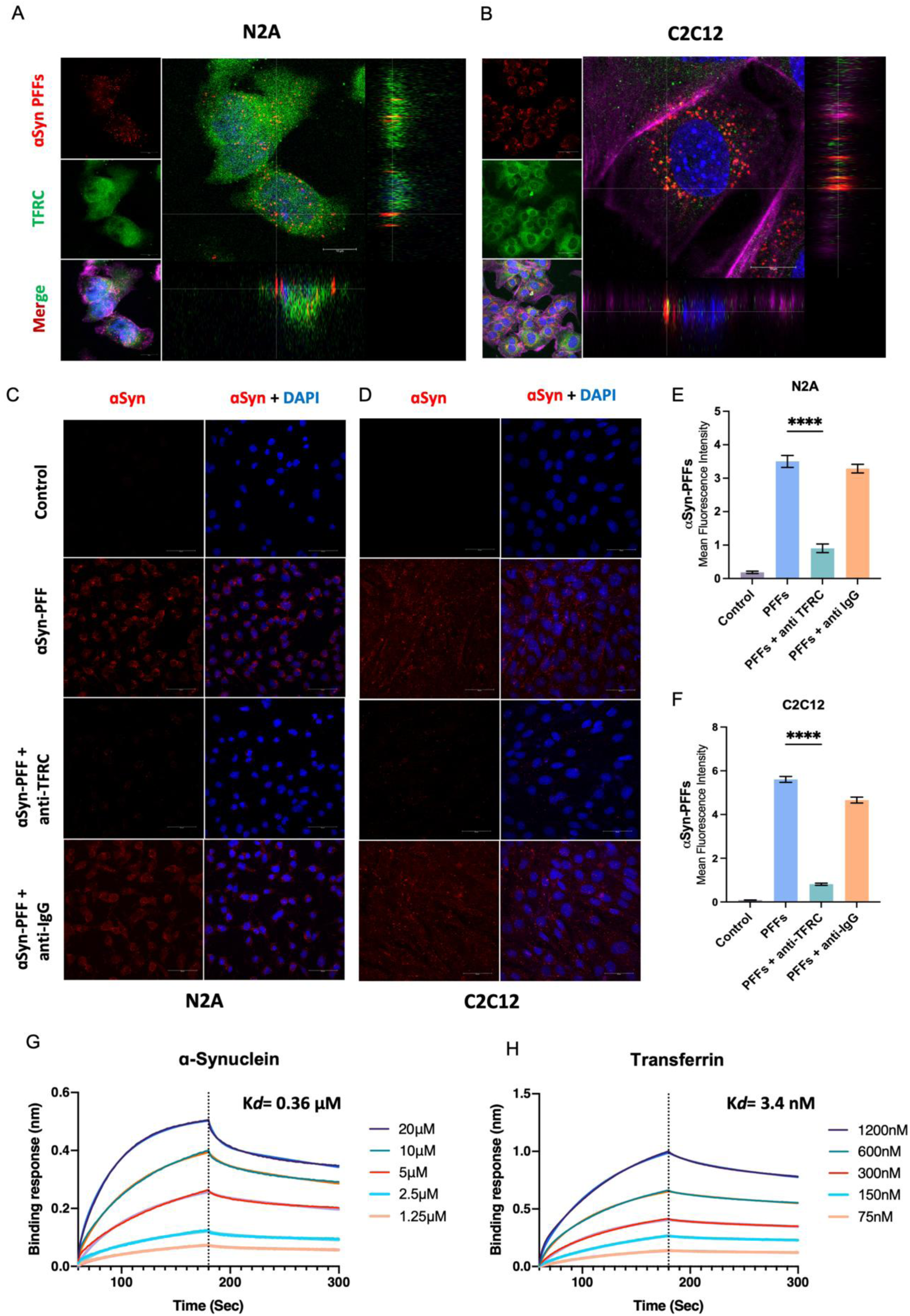
α-Synuclein interacts with TFRC. A, B. Representative confocal micrographs and orthogonal sections of N2A (A) and C2C12 cells (B) labeled for αSyn-PFFs (red), TFRC (green), F-actin (magenta), and DAPI (blue) depicting αSyn-TFRC colocalization. C, D. Representative confocal micrographs of N2A (C) and C2C12 cells (D) labeled for αSyn-PFFs (red), and DAPI (blue). E, F. Quantification of the mean fluorescence intensity of αSyn-PFFs at the surface in N2A (E) and C2C12 cells (F). G, H. Biolayer Interferometry (BLI) sensograms showing interactions between TFRC and αSyn (G) and Transferrin (H) (n = 3 independent experiments). Data is shown as mean ± sem, n = 3 independent experiments. One-way ANOVA followed by Tukey’s correction, \*\*\*\**P* < 0.0001.

### 8. αSyn promotes TFRC-dependent iron accumulation and lipid peroxidation in neuronal and muscle cells

To further determine whether αSyn induces a ferroptosis-associated phenotype, we examined whether the αSyn–dependent increase in surface Transferrin receptor 1 (TFRC) observed in our surface proteomics analysis leads to functional alterations in iron metabolism and lipid peroxidation. First, we validated TFRC expression in N2A and C2C12 cells by western blot analysis. Cells were treated with αSyn-PFFs alone or after pre-incubation with an anti-TFRC antibody, with anti-IgG serving as an isotype control. Consistent with our surface proteomics results, αSyn-PFFs treatment significantly increased TFRC protein levels in both N2A and C2C12 cells (Figure S5A-C). Notably, this increase was abolished in cells pre-treated with the TFRC-blocking antibody, indicating that αSyn interaction with TFRC contributes to the observed increase in receptor expression. As TFRC is a key mediator of cellular iron uptake, we next assessed whether αSyn-induced TFRC upregulation results in increased intracellular iron levels. Cells treated with αSyn-PFFs were stained with FerroOrange, a fluorescent probe for intracellular ferrous iron (Fe²⁺) and confocal microscopy was performed (Figure 9A-D). αSyn-PFFs treatment significantly increased FerroOrange fluorescence intensity in both N2A and C2C12 cells, indicating elevated intracellular iron accumulation. We then examined whether this iron accumulation promotes lipid peroxidation, a hallmark of ferroptotic cell death [29]. N2A and C2C12 cells were treated with erastin, a canonical ferroptosis inducer, or with αSyn-PFFs in the presence or absence of ferrostatin-1, a potent inhibitor of lipid peroxidation and ferroptosis[76]. Lipid peroxidation was first assessed by immunostaining for 4-HNE (Figure 9E-H). αSyn-PFFs treatment significantly increased 4-HNE levels in both cell types, comparable to erastin in N2A cells and even more pronounced in C2C12 cells. Importantly, pre-treatment with ferrostatin-1 significantly reduced αSyn–induced 4-HNE accumulation. To further quantify lipid peroxidation, we performed the BODIPY-C11 lipid oxidation assay, a ratiometric probe that shifts fluorescence from red to green upon oxidation (Figure 9I, J). Consistent with the 4-HNE results, αSyn-PFF treatment significantly increased BODIPY-C11 oxidation in both N2A and C2C12 cells, indicating elevated lipid peroxidation. This effect was comparable to erastin treatment and was significantly attenuated by ferrostatin-1, confirming lipid peroxidation. Together, these findings demonstrate that αSyn promotes TFRC-dependent iron accumulation and lipid peroxidation in both neuronal and muscle cells, establishing a mechanistic link between αSyn–TFRC interaction and ferroptosis-associated oxidative damage. These results are consistent with our in vivo observations in A53T mice, in which increased TFRC expression, iron accumulation, and lipid peroxidation were detected in both brain and skeletal muscle.

**Figure 9.**
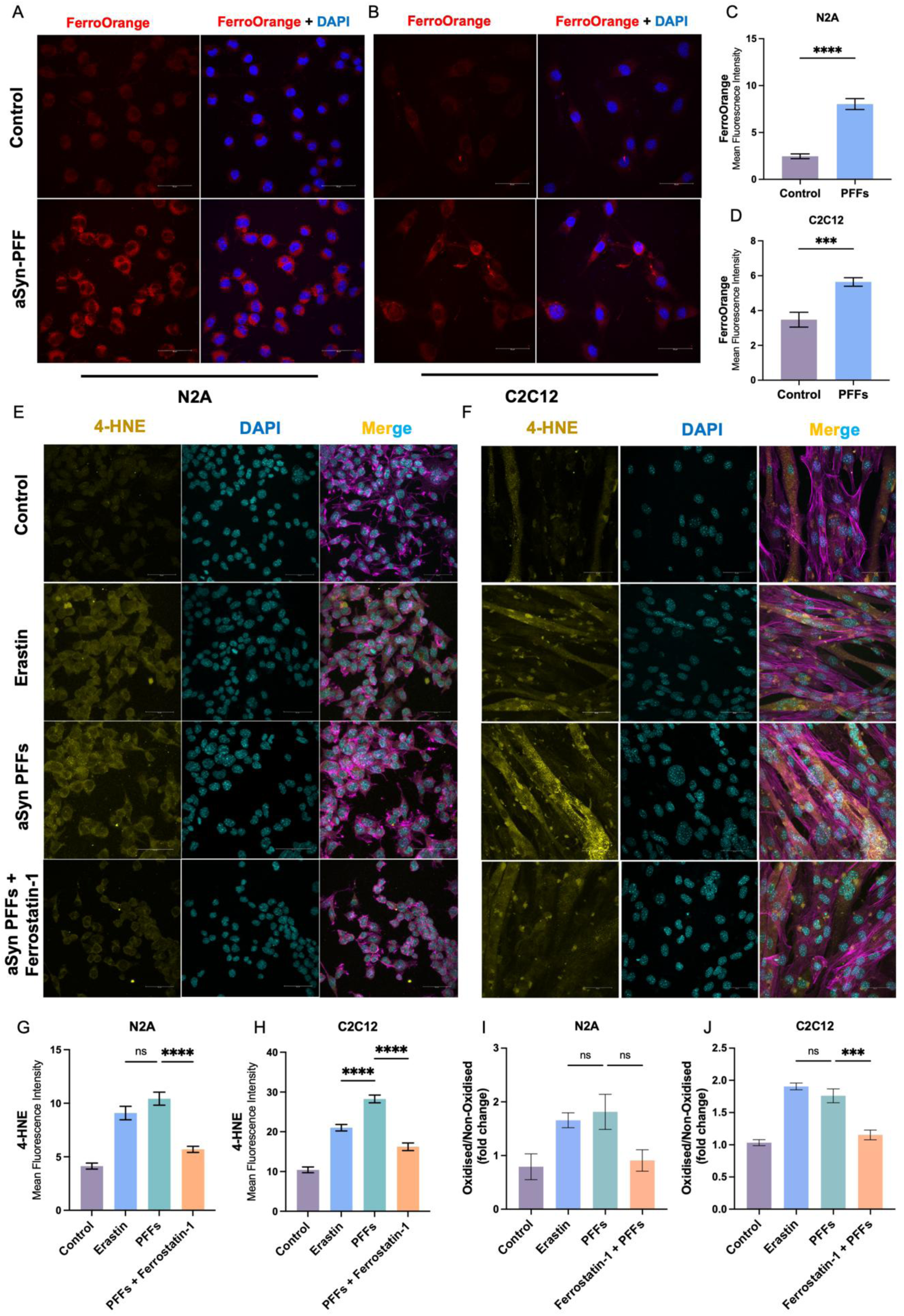
α-Synuclein induces TFRC-dependent ferroptosis. A, B. Representative confocal micrographs of N2A (A) and C2C12 cells (B) labeled for intracellular iron using FerroOrange (red) and DAPI (blue). C, D. Quantification of the mean fluorescence intensity of FerroOrange in N2A (C) and C2C12 cells (D). E, F. Representative confocal micrographs of N2A (E) and C2C12 cells (F) labeled for 4-HNE (yellow), F-actin (magenta), and DAPI (cyan). G, H. Quantification of the mean fluorescence intensity of 4-HNE in N2A (G) and C2C12 cells (H). I, J. Quantification of the fold change in oxidised/non-oxidised ratio of BODIPY-C11 in N2A (I) and C2C12 cells (J) Data is shown as mean ± sem, n = 3-5 independent experiments. One-way ANOVA followed by Tukey’s correction, \*\*\*\**P* < 0.0001 and \*\*\**P* < 0.001.

### 9. Plasma proteomic analysis reveals systemic metabolic alterations associated with ferroptosis in aged PD mice

To determine whether ferroptosis-associated molecular alterations observed in the brain and the skeletal muscle are reflected at the systemic level, we performed targeted multiple reaction monitoring (MRM) analysis of circulating plasma proteins from 18-month-old A53T mice. MRM analysis revealed significant dysregulation of several metabolic and stress-associated factors known to regulate oxidative stress, lipid metabolism, and inflammatory signaling (Figure 10A-I). Among the proteins, the adipokine leptin (LEP), the metabolic peptide apelin (APLN), the ER stress chaperones BiP (HSPA5), calreticulin (CALR), and the muscle catabolic regulator myostatin (MSTN) were elevated in the plasma of A53T mice (Figure 10A-E). Leptin elevation acts as a pro-inflammatory adipokine, driving reactive oxygen species (ROS) production and lipid oxidation. Leptin also activates the JAK/STAT and PI3K pathways, increasing iron uptake. Chronic high leptin levels impair mitochondrial function, promoting lipid peroxidation and increasing ferroptosis vulnerability[77,78]. Apelin modulates metabolism and mitochondrial health and counters lipid peroxidation by activating AMPK signaling[79]. Thus, increased apelin levels suggest compensatory mechanisms in response to ferroptosis. Bip is an ER stress chaperone involved in the unfolded protein response, counteracting proteotoxic and oxidative stress [80,81], whereas calreticulin is an ER chaperone regulating calcium homeostasis and protein folding. Calreticulin upregulation has been shown to induce calcium dysregulation, thus promoting mitochondrial calcium uptake and inducing ROS generation and lipid peroxidation, linking ER stress, a well-established feature of αSyn toxicity, to the broader ferroptotic oxidative cascade[82,83]. Myostatin inhibits muscle hypertrophy and repair pathways while activating catabolic pathways, mitochondrial impairment, and ROS production[84,85]. Thus, increased myostatin levels suggest elevated lipid peroxidation in muscle cells, thereby increasing ferroptosis sensitivity, consistent with the neuromuscular alterations observed in the A53T mice. Conversely, levels of several circulating proteins with protective antioxidant and metabolic functions were significantly reduced, including neuropeptide Y (NPY), adiponectin (ADIPOQ), apolipoprotein H (APOH), and fibroblast growth factor 21 (FGF21) (Figure 10G-I). NPY is a neuroprotective peptide against microglial activation, calcium excitotoxicity, and nitric oxide-mediated mitochondrial damage[86]. Thus, downregulated NPY removes a protective mechanism against ROS accumulation and ferroptotic damage. Adiponectin is a cytokine secreted by adipocytes that regulates lipid metabolism and glucose uptake. Adiponectin has been recently shown to confer protection against neuroinflammation and ferroptosis through AMPK signalling [87,88]. Apolipoprotein H functions to transport lipids and clear oxidized lipids [89]. Therefore, downregulation of APOH increases the risk of ferroptosis. Similarly, Fibroblast Growth Factor 21 (FGF21) is a metabolic stress hormone and a myokine involved in mitochondrial function, energy homeostasis, NMJ maintenance, and muscle integrity. FGF21 boosts NRF2 signalling, glutathione synthesis, and limits mitochondrial ROS generation[90–92]. FGF21 has recently been shown to enhance survival in Amyotrophic Lateral Sclerosis (ALS), a neurodegenerative disease characterized by progressive muscle weakness[93]. The reduced FGF21 levels therefore suggest impaired metabolic adaptation and systemic stress signaling in the A53T mice, further supporting the brain and muscle data showing downregulation of the NRF2 response and NMJ degeneration. Therefore, alterations in these key plasma proteins suggest compromised systemic antioxidant defenses and lipid detoxification pathways, thereby facilitating lipid peroxidation and ferroptotic susceptibility in the A53T mice. The plasma MRM profile supports the existence of a systemic ferroptosis-associated state. Importantly, these plasma alterations also complement our previous MRM findings, which demonstrated systemic inflammatory activation in A53T mice. In the context of our findings showing increased TFRC expression and ferroptosis phenotype in both brain and skeletal muscle of the A53T mice, the plasma MRM profile supports the existence of a systemic ferroptosis-associated state. These results also suggest that αSyn–driven TFRC-mediated ferroptosis may not be restricted to individual tissues but instead contributes to a broader systemic pathology linking brain and muscle degeneration in the A53T mouse model of PD.

**Figure 10.**
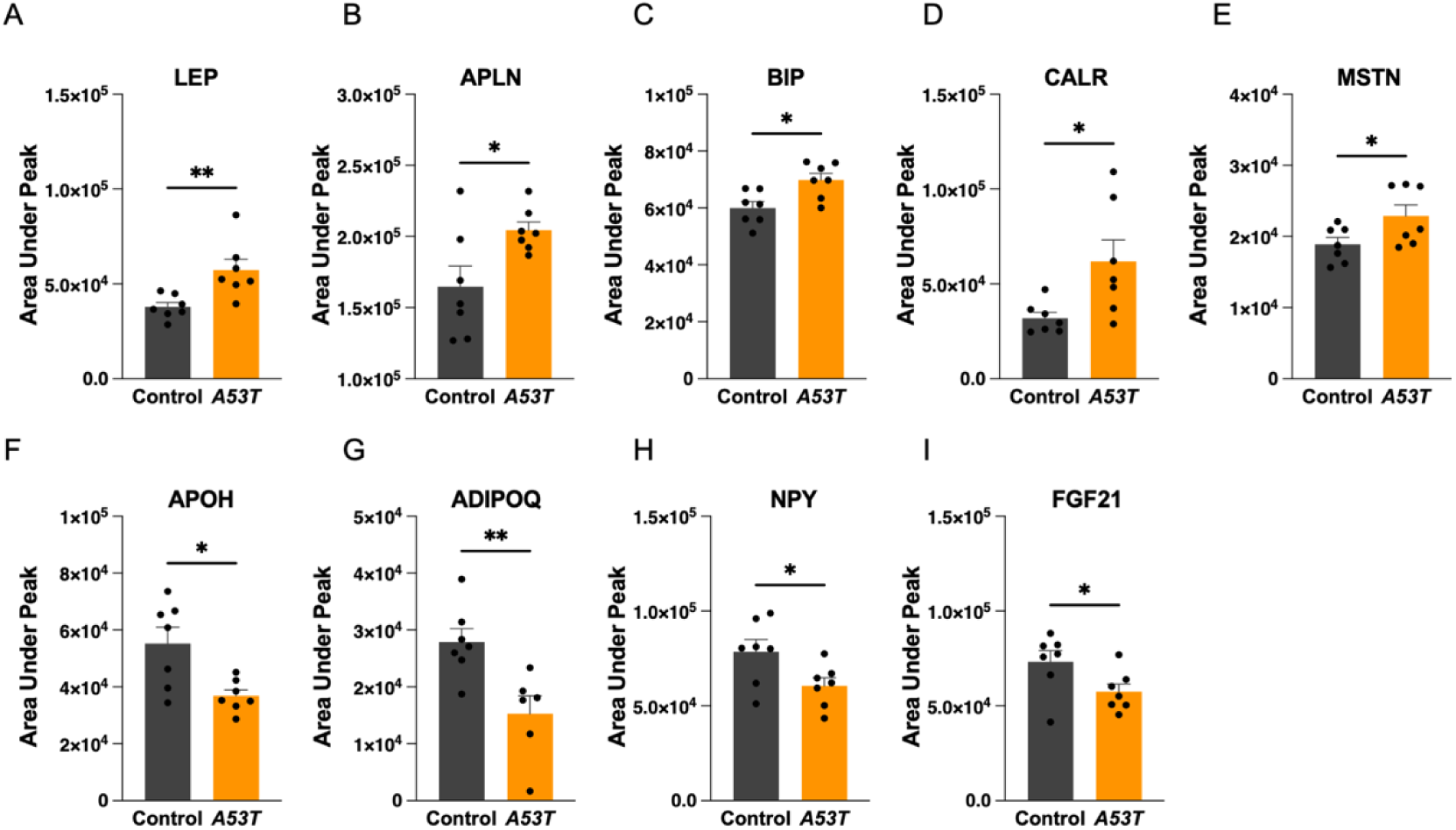
Plasma MRM-MS reveals systemic ferroptosis signature in A53T mice. A-I. MRM based relative quantification of plasma protein, namely (A) LEP, (B) APLN, (C) BIP, (D) CALR, (E) Myostatin, (F) Apolipoprotein H, (G) Adiponectin Q, (H) Neuropeptide Y, and (I) Fetal Growth Factor 21 represented as the area under the peak in control and A53T mice. Data presented as mean ± SEM; an unpaired two-tailed Student’s t-test was employed, with \*\**P* < 0.01 and \**P* < 0.05.

## Discussion

Parkinson’s disease (PD) has historically been conceptualized as a brain-restricted disorder, defined by the progressive loss of dopaminergic neurons in the substantia nigra and the accumulation of misfolded αSynuclein in Lewy bodies[2,3]. However, a growing body of evidence positions PD as a multisystem disorder in which peripheral tissues are also involved [5,18]. Although the CNS is primarily affected in PD, the other organ that shows pronounced symptoms is the skeletal muscle. PD motor symptoms, such as bradykinesia, tremors, and postural instability, directly involve the skeletal muscle [40,41]. PD patients often exhibit approximately 30-50% increase in the incidence of severe muscle wasting or sarcopenia, frailty, and muscle dysfunction, which exacerbates disease progression and reduces quality of life[40,94]. However, the molecular links between the brain and muscle pathology remain poorly explored, and studies linking the brain and skeletal muscle αSyn pathology in PD are scarce. Using a transgenic mouse model of PD expressing mutant human A53T αSyn, we investigate αSyn aggregation-associated pathology in both brain and skeletal muscle. A53T mice exhibit classical PD symptoms, including reduced motor coordination on the rotarod and pole tests and increased activity in the CLAMS, indicating motor impairments and behavioral changes. A53T mice also show reduced body weight compared to controls. A53T mutations in αSyn accelerate aggregation and promote neurotoxicity[95]. Consistent with previous studies, at the molecular level, we found that αSyn and phosphorylated αSyn (pS129) levels were increased in the brain, indicating αSyn accumulation. Tyrosine hydroxylase (TH), a marker of dopaminergic neurons, was downregulated, and decreased TH-positive neurons were observed in the substantia nigra region of A53T mice, indicating dopaminergic neuronal degeneration due to αSyn aggregation. The A53T mice exhibited decreased muscle mass in all four representative muscles, with muscle atrophy and reduced function. The gastrocnemius muscle of A53T mice showed reduced cross-sectional area along with an increase in the proportion of small fibers and a reduction in large fibers. Fast myofibers, which are glycolytic, are larger than slow, type I oxidative muscle fibers[96]. This suggests that the A53T mice have an increased proportion of slow myofibers as previously reported in PD patients[97–100]. Each myofiber type possesses unique contractile properties due to the expression of different myosin heavy chain (MyHC) isoforms[101]. The fiber-type alterations, therefore, may contribute to the observed impairment of grip strength in the A53T mice. The αSyn and pS129 levels were also elevated in the gastrocnemius muscle, suggesting the role of αSyn in the observed muscle alterations.

The neuromuscular junction (NMJ) represents the critical interface between the motor neuron and skeletal muscle, and its structural and functional integrity is essential for voluntary movement[102]. Interestingly, we found elevated αSyn and pS129 expression specifically at the NMJs in the A53T mice. This was accompanied by NMJ-associated defects, including fragmentation and an increased number of nuclei. αSyn is a presynaptic protein with a role in synaptic vesicle trafficking and neurotransmitter release[7,103]. Thus, pathological αSyn accumulation at the NMJs is expected to disrupt synaptic signaling. The molecular analysis of NMJ-associated proteins reveals downregulation of NCAM1, LRP4, and MuSK and upregulation of AChR, suggesting altered synaptic structure and failure in reinnervation and regeneration. These molecular changes were accompanied by reduced ex-vivo muscle contractility, suggesting that pathological αSyn accumulation disrupts NMJ structure and signaling, contributing to impaired muscle function. The NMJ degeneration has been previously reported in other motor neuron disease models and in aged muscle[104–108]. This further explains the muscle atrophy and functional deficits observed in A53T mice. These results also confirm that the A53T mouse model recapitulates essential pathological features of PD and provides a robust system to investigate mechanisms linking αSyn pathology in the brain and the skeletal muscle.

Our plasma MRM-MS profiling of A53T mice reveals a robust systemic inflammatory signature that mirrors and extends beyond the CNS pathology. The elevation of TNF, IL-1β, IL-18, C3, GFAP, and CX3CL1 in circulation reflects an active innate immune system and cytokine release. More importantly, elevated circulating complement C3 levels suggest that innate immune amplification in A53T mice extends to the lectin and alternative complement cascades. Complement C3 has previously been implicated in NMJ degeneration and synapse elimination in Myasthenia gravis[109], as also observed in our A53T model. Detection of elevated GFAP and CX3CL1 levels in plasma indicates compromised blood-brain barrier integrity and active neuroinflammation, allowing CNS-derived proteins to enter the systemic circulation. Together, the plasma inflammatory profile indicates that A53T mice exhibit a systemic neuroimmune activation state, likely contributed and further amplified by tissue-level degeneration in both brain and skeletal muscle. The coexistence of brain degeneration and skeletal muscle pathology, along with systemic inflammation, raises the possibility of shared pathological communication between these tissues.

Our proteomic analyses of brain and skeletal muscle tissues from 18-month-old A53T and control mice further support the existence of brain–muscle crosstalk, revealing shared molecular pathways involved in both tissues. The brain-specific enrichment of synaptic vesicle cycling and neuroactive ligand-receptor pathways corroborates our phenotypic observations of dopaminergic neurodegeneration, while muscle-specific enrichment of HIF-1 signalling and mitochondrial organization pathways reflects the metabolic reprogramming characteristic of atrophied and denervated muscle[110,111]. Notably, dysregulation of the neuromuscular pathway in the muscle proteome provides corroborating evidence for our NMJ findings and suggests that muscle defects involve disruption of intrinsic neuromuscular signalling.

Among the shared pathways, the identification of iron uptake and transport in the brain and ferroptosis in the muscle represents the central and novel finding of our proteomic analysis in A53T mice. Ferroptosis is a regulated form of cell death characterized by iron accumulation and lipid peroxidation[29]. Our proteomic analyses identified numerous ferroptosis-associated proteins, including markers, drivers, and suppressors, in both tissues. Our biochemical validation confirmed increased iron accumulation, elevated lipid peroxidation, and suppression of key antioxidant defenses. Specifically, increased expression of the iron uptake receptor TFRC, together with decreased expression of ferroptosis-protective proteins such as SLC7A11 and GPX4, indicates increased ferroptotic susceptibility in both brain and skeletal muscle of A53T mice. The downregulation of SLC7A11 impairs cystine import and thus glutathione (GSH) biosynthesis. GPX4 utilizes GSH to reduce membrane-embedded phospholipid hydroperoxides[57,60]. Thus, concurrent downregulation of GPX4 and SLC7A11 abolishes the primary enzymatic safeguard against ferroptotic lipid peroxidation and cell death[31,112]. Interestingly, brain and skeletal muscle exhibited distinct patterns of iron homeostasis, despite elevated TFRC levels in both tissues. FTH1 levels were reduced in the skeletal muscle of A53T mice compared to the brain. FTH1 encodes the heavy subunit of ferritin, the primary intracellular iron storage protein. FTH1, through its ferroxidase activity, oxidizes ferrous iron (Fe²⁺) to the ferric form (Fe³⁺) for safe, soluble storage, preventing toxicity from free iron [113,114]. The elevated brain FTH1 likely reflects a compensatory but insufficient response to iron overload, whereas reduced skeletal muscle FTH1 suggests that muscle cells are not only importing more iron but are also failing to sequester it effectively, most likely due to ferritinophagy-mediated ferritin turnover or impaired ferritin biosynthesis. Thus, suggests a larger, unsequestered labile iron pool (LIP) in the muscle, rendering it more susceptible to iron-induced oxidative stress. Importantly, although ferroptosis has been extensively studied in the PD brain[26,27,115], its role in the skeletal muscle has not been reported previously. Our findings, therefore, identify ferroptosis as a previously unrecognized mechanism contributing to muscle pathology in PD.

The identification of shared NRF2 and associated antioxidant protein suppression in both brain and skeletal muscle of A53T mice further suggests a failure of antioxidant defense across tissues. NRF2 is the master transcriptional regulator of the cellular antioxidant response and a known modulator of ferroptosis susceptibility, controlling the expression of SLC7A11, GPX4, GPX1, ferritin subunits, catalase, and peroxiredoxins, among others[30,63,116]. Thus, iron overload, combined with antioxidant defense failure, leads to unrestrained lipid peroxidation, as evidenced by elevated 4-HNE levels in both the brain and skeletal muscle of the A53T mice.

Mechanistically, our study reveals a potential upstream molecular determinant linking αSyn pathology to ferroptotic signaling. Our cell-surface proteomic analysis following αSyn-PFF treatment of N2A neuroblastoma and C2C12 myoblast cells provides detailed information on αSyn-induced cell-surface alterations and identifies TFRC as a novel cell-surface protein, significantly upregulated in both cell types. Through microscopic and biophysical interaction studies, we identified a direct interaction between αSyn and TFRC. Transferrin receptor (TFRC) has been established as a ferroptosis marker that facilitates intracellular iron uptake and drives iron-induced lipid peroxidation[71,72]. Blocking TFRC reduced αSyn binding at the cell surface, and functional assays demonstrated that αSyn exposure increased intracellular iron levels and lipid peroxidation in both neuronal and muscle cells. These findings suggest that TFRC may serve as a receptor or facilitator for αSyn-mediated iron dysregulation. By promoting iron influx, the αSyn-TFRC interaction may initiate a cascade of events that ultimately drives ferroptotic cell death.

In addition to tissue-level changes, our plasma MRM-MS analysis revealed systemic metabolic and inflammatory alterations consistent with ferroptosis-associated stress. The levels of circulating pro-inflammatory and metabolic stress mediators, such as Leptin, Apelin, and ER stress proteins, including Bip and Calreticulin, were elevated in the A53T mice. Conversely, protective metabolic regulators such as Adiponectin, NPY, and FGF21 were significantly reduced. These systemic metabolic alterations complement our earlier observations of circulating inflammatory cytokines and collectively indicate that PD pathology in the A53T mice is accompanied by a bidirectional relationship between inflammation and ferroptosis. As NF-κB activation in PD increases inflammatory cytokines such as TNF and IL-1β, and suppresses NRF2 and ultimately SLC7A11 expression[117–120], and in turn, ferroptotic cell death releases DAMPs that further activate innate immune responses[121,122]. This further suggests that these two plasma signatures are not independent but rather mutually existing components of a systemic pathology that mediates neurodegeneration and muscle atrophy in the A53T mice.

Taken together, our results suggest that αSyn-driven iron dysregulation and ferroptosis contribute to shared pathology in both brain and muscle and further extend systemically. αSyn interaction with TFRC promotes iron accumulation and lipid peroxidation, while suppression of antioxidant pathways further increases ferroptotic vulnerability. These processes are accompanied by systemic metabolic and inflammatory changes detectable in circulation. Thus, our study highlights a previously unrecognized αSyn–TFRC–ferroptosis axis linking neurodegeneration and muscle dysfunction in PD.

In conclusion, the present study provides the first comprehensive evidence that αSyn pathology drives ferroptosis as a shared mechanism across the brain-muscle axis in PD. This process is mediated through direct αSyn-TFRC interaction. These findings also present the first evidence of αSyn-driven ferroptosis in the skeletal muscle in PD. Our findings expand current understanding of PD from a brain-centric disorder to a systemic disorder, suggesting that targeting iron metabolism, TFRC signaling, or ferroptosis-related pathways may represent promising therapeutic strategies for mitigating both neuronal and peripheral manifestations of PD.

## Materials and Methods

### 1. Animals and Study Design

B6;C3-Tg(Prnp-SNCA*A53T)83Vle/J mice were purchased from The Jackson Laboratory, USA, and genotyped accordingly. Heterozygous male mice carrying one A53T αSyn allele were used in the study. Mice were housed in standard conditions under a 12-hour light/dark cycle and monitored for the development of PD symptoms. Animals were euthanized after disease onset at 18 months of age, and tissues such as brain, skeletal muscle, and plasma were collected. All the animal experiments and procedures were approved by the Institutional Animal Ethics Committee (IAEC), protocol number RCB/IAEC/2022/141.

### 2. Behavioral Experiments

#### Rotarod

Mice were placed on an accelerating rotarod (Harvard Apparatus), with speed increasing from 0 to 50 rpm, and the latency to fall was recorded. Mice were trained for 3 days before the final experiment. 3 trials were performed with an interval of 15 minutes each, and the data are presented as a mean latency time on the rotarod.

#### Pole test

Used to measure motor coordination in mice. The mice were placed on a vertical pole wrapped in banded gauze (9 mm diameter and 75 cm in height) with their heads oriented upwards. The average time to turn and reach the bottom of the pole was recorded. The mice were trained for 3 days, 3 times each, before the final experiment.

#### Grip strength

To assess the neuromuscular strength of the mice, grip strength was tested (GT3, Panlab, Harvard Apparatus). The mice were placed on a metal grid and allowed to grasp using both fore and hind limbs. The mouse was gently pulled from the tail, and the maximum holding force was recorded. Three measurements per mouse were recorded and are presented as mean grip strength normalised to the mouse’s body weight.

### 3. Muscle Contractility

The muscle contractile properties of non-transgenic (control) and human αSyn hemizygous (A53T^+/-^) transgenic mice were measured at 18 months of age using a muscle contractility measurement apparatus (iWORX, IX-TA-220 data recorder and FT-302 force transducer) as previously reported [123]. Briefly, the control and A53T mice were anesthetized, and the gastrocnemius muscle was partially excised by removing it from the distal tendon while keeping the proximal tendon intact. The isolated GAS muscle was attached to a force transducer and a recording system to assess its contractile properties. Electrical pulses ranging from 25 Hz to 300 Hz were applied using a pair of electrodes, and the resulting force–frequency relationship was recorded. The generated forces were plotted as absolute force and maximum force in grams.

### 4. Western blotting

Brain and muscle tissue were harvested from 18-month-old A53T mice and flash frozen in liquid nitrogen. The tissues were homogenized in ice-cold radioimmunoprecipitation assay (RIPA) buffer (Sigma, R0278) containing protease/phosphatase inhibitor cocktail (Thermo Scientific, 78446), followed by intermittent vortex mixing for 30 min on ice. The lysate was centrifuged at 16,000 g for 15 min at 4°C. The supernatant was collected, and the protein concentration was measured using the Pierce BCA Protein Assay kit (Thermo Scientific; 23225). The protein samples were separated using SDS-PAGE and transferred to a 0.45 µm PVDF membrane using standard wet transfer protocols. Upon transfer, the membranes were blocked with 5% skimmed milk or 5% BSA in Tris-buffered saline with 0.01% Tween 20. Primary antibody incubation was performed in blocking buffer at 4°C overnight, and subsequent HRP-conjugated secondary antibody incubation was done at room temperature for 1 hr. Blots were developed using the HRP substrate (Millipore; WBLUF0100) and imaged on ImageQuant LAS 4000 (GE Healthcare). The densitometric analysis was performed using ImageJ software.

For cell culture experiments, cells upon treatment were washed 3 times with 1X Phosphate-buffered saline and collected after treatment. The cells were lysed in RIPA buffer with probe sonication for 10 seconds. The lysate was centrifuged at 16,000 g for 15 min at 4°C. The supernatant was collected, and the protein concentration was measured, followed by procedure similar to that discussed for western blotting.

### 5. Iron Measurement

50 mg of whole brain and muscle tissue were homogenized in 0.9% NaCl solution and probe sonicated at 25% amplitude for 10 sec on ice. Later, the lysate was centrifuged at 10,000g for 20min at 4°C, and the supernatant was collected. Iron measurement was performed as per the manufacturer’s instructions (Sigma-Aldrich, MAK472). The samples were incubated with reagents for 40 minutes at room temperature. The optical density was measured at 590 nm. The concentration was calculated using the iron standard curve. Each sample was run in duplicate.

### 6. Mass Spectrometry (MS)

#### Sample preparation

1 mg of whole brain and gastrocnemius muscle tissue was homogenized on ice. The homogenate was added to 500 μL RIPA buffer (Sigma, R0278) containing a 1X halt protease/phosphatase inhibitor cocktail (Thermo Scientific, 78446), and the mixture was probe-sonicated for 10sec at 25% amplitude. The lysate was centrifuged at 16,000g for 30 min at 4°C. The supernatant was collected, and the protein concentration was measured using the Pierce BCA Protein Assay kit (Thermo Scientific, 23225). Further, 500 µg of protein was collected, and ice-cold acetone was added at the (1:5) protein to acetone ratio. The mixture was incubated at -20°C overnight and later centrifuged at 8000g for 30min at 4°C. The precipitate was washed with ice-cold acetone. Later, the precipitate was dissolved by adding 8M urea. The solubilized protein concentration was estimated, and 100 μg protein was further reduced with 10 mM dithiothreitol (DTT) at 56°C for 30 min and alkylated with 20 mM iodoacetamide (IAA) at room temperature for 1 h. The proteins were digested with mass spectrometry-grade trypsin (1:15 *w/w*, Pierce, Thermo Fisher Scientific, Waltham, MA, USA) at 37 °C for 18 h. The digested peptide samples were next desalted using Oasis® HLB 1cc (10mg) extraction cartridges (Waters, Milford, MA, USA). The peptides were eluted using 200μl of elution reagent (70% acetonitrile and 30% water (v/v) with 0.1% formic acid). Eluted peptides were vacuum-dried and stored at -80°C until further use.

#### Data-independent acquisition and processing

The vacuum-dried desalted tissue samples were resuspended in solvent A (98% H2O and 2% acetonitrile (v/v) with 0.1% formic acid) and injected into the ZenoTOF 7600 mass spectrometer (SCIEX, MA, USA) coupled with ACQUITY UPLC M-Class system (Waters, MA, USA). 1µg digest from each sample was loaded, in triplicate, onto a Luna Micro Trap (20 x 0.3 mm, 5 µm, 100 Å, Phenomenex, Torrance, CA, USA) column at a flow rate of 5 μL/min for 3 min. The tryptic peptides were then eluted from nanoEasseTM M/Z HSS T3 (100 Å, 1.8 µm, 300 µm x 150 mm, Waters, MA, USA) analytical column at a flow rate of 5 μL/min in a linear gradient of 5% solvent B (100% ACN (v/v) with 0.1% formic acid) to 80% solvent B in 45 min with a total run time of 50 min. A SWATH-DIA (Sequential Windowed Acquisition of All Theoretical Fragment Ion Mass Spectra- Data-Independent Acquisition) acquisition scheme with 65 overlapping variable windows spanning a mass range of 350-1250 Da and an accumulation time of 20 ms per window was employed.

The acquired DIA-MS data were analysed on Spectronaut Pulsar (v18.2.23, Biognosys, Schlieren, Zurich, Switzerland). A de novo spectral library was first prepared by searching the entire data set against the isoform included UniProtKB *Mus Musculus* proteome database (UP000000589, 21,856 gene count) with default search parameters. The brain data library comprised 33,001 precursors and 25,354 peptides corresponding to 3,424 protein groups, whereas the muscle data library comprised 26,757 precursors and 19,141 peptides corresponding to 2297 protein groups. The raw files were then searched against the generated library and the mouse database using default settings: quantification at the MS2 level, differential abundance testing using an unpaired Student’s t-test, and global TIC was used for data normalisation. FDR was set to 1% at both peptide and protein levels. The extracted areas under the peptide peaks were further used for quantitative analysis of the plasma proteome.

#### Multiple reaction monitoring (MRM)

The stored plasma samples were diluted twenty-fold in 100 mM ammonium bicarbonate (ABC), and 20 μL of the diluted neat plasma was aliquoted from each sample for further processing. Diluted plasma was further reduced with 10 mM dithiothreitol (DTT) at 56° C for 30 min and alkylated with 20 mM iodoacetamide (IAA), at room temperature (RT) for 1 h. The proteins were digested with mass spectrometry-grade trypsin (1:15 w/w, Pierce, Thermo Fisher Scientific, Waltham, MA, USA) at 55 °C for 6 h. The digested peptide samples were next desalted using Oasis® HLB 1cc (10mg) extraction cartridges (Waters, Milford, MA, USA). The peptides were eluted using 200μl of elution reagent (70% acetonitrile and 30% water (v/v) with 0.1% formic acid). Eluted peptides were vacuum-dried and stored at -80°C until further use.

Targeted mass spectrometry was carried out in MRM mode on the Triple Quad 6500+ system (SCIEX, MA, USA) coupled with ExionLC (SCIEX, MA, USA). For this, the y and b ions of the target proteins were selected from the SRMatlas Database [124]. Initially, all the ions were scanned in an unscheduled MRM mode on a pooled plasma sample to optimise the retention time (RT) of the ions. The product ions with Gaussian peak shapes and intensities greater than 100 counts per second (cps) were further selected and acquired in a scheduled MRM mode on the pooled sample. Finally, the optimised method consisting of 253 transitions with a target scan time of 1.3 s, MRM detection window of 120 s and total run time of 70 min, was used to acquire data for all the plasma samples. The samples were eluted at a flow rate of 0.2 mL/min from an Acquity UPLC CSH TM C18 analytical column (1.7 µm, 2.1 x 150 mm, Waters, MA, USA) in a linear gradient of 2% solvent B (100% ACN (v/v) with 0.1% formic acid) to 95% solvent B in 65 min with a total run time of 70 min. The raw MRM data were analyzed using the Analytics software integrated with SCIEX OS platform (v 3.1.6.44, SCIEX, MA, USA). The extracted peaks were selected based on expected RT, pre-processed by smoothing on a moving average, and filtered based on the following criteria: minimum peak height of 100 cps and minimum signal/noise ratio of 2. All selected spectral peaks were checked manually to ensure correct peak detection and accurate integration. The areas under the peak of selected product ions were used for all downstream processing.

#### Surface Proteome

N2A and C2C12 cells were treated with 2.5 μM of ɑSynuclein pre-formed fibrils and incubated for 2 h at 37°C. Upon incubation, cells were washed 3 times with 1X Phosphate-buffered saline to remove media components, then collected by gentle scraping. The surface protein biotinylation reagent EZ-Link™ Sulfo-NHS-LC Biotin (ThermoFisher, 21335) was added to the cell suspension at 1.0 mg/mL and incubated for 30 min at 4°C in end-to-end rotation. The labelling was quenched by washing the cell suspension with 100 mM glycine three times. After labelling was complete, cells were lysed in RIPA buffer containing protease/phosphatase inhibitor cocktail. The protein concentration was estimated, and 1 mg protein was acetone-precipitated by adding ice-cold acetone at a 1:5 protein-to-acetone ratio. The solution was incubated at -20°C overnight and centrifuged at 8000*g* for 30min at 4°C. The protein precipitate was washed with ice-cold acetone and solubilized by adding 8 M urea. The solubilized protein concentration was estimated, and 500 μg protein was further reduced with 10 mM dithiothreitol (DTT) at 56°C for 30 min and alkylated with 20 mM iodoacetamide (IAA) at room temperature for 1 h. The proteins were digested with MS-grade trypsin (1:15 *w/w*, Pierce, Thermo Fisher Scientific, Waltham, MA, USA) at 37 °C for 18 h. The digested peptide samples were next desalted using Oasis® HLB 1cc (10mg) extraction cartridges (Waters, Milford, MA, USA). The peptides were eluted with 200 μL of 70% acetonitrile in water (v/v) containing 0.1% formic acid and vacuum-dried. The peptides were dissolved in sodium phosphate buffer, and immunoprecipitation was performed using streptavidin beads to enrich biotinylated peptides. The biotinylated peptides were eluted from the streptavidin beads and vacuum dried.

The vacuum-dried desalted samples were resuspended in solvent A (98% H2O and 2% acetonitrile (v/v) with 0.1% formic acid) and then injected into the ZenoTOF 7600 mass spectrometer (SCIEX, MA, USA) coupled with ACQUITY UPLC M-Class system (Waters, MA, USA). An equal amount of digest from each sample was loaded, in triplicate, onto a Luna Micro Trap (20 x 0.3 mm, 5 µm, 100 Å, Phenomenex, Torrance, CA, USA) column at a flow rate of 5 μL/min for 3 min. The tryptic peptides were then eluted from nanoEasseTM M/Z HSS T3 (100 Å, 1.8 µm, 300 µm x 150 mm, Waters, MA, USA) analytical column at a flow rate of 5 μL/min in a linear gradient of 5% solvent B (100% ACN (v/v) with 0.1% formic acid) to 80% solvent B in 45 min with a total run time of 50 min. A DDA acquisition scheme was employed with a dynamic exclusion window of 5 sec for MS1 scans (m/z 300-1500 Da), from which the top 45 ions were selected for MS2 scans (200-1800 m/z). The collision energy was set as dynamic, while a declustering potential of 80 V is used. The acquired raw files were converted to mzML file using MSConvertGUI for Fragpipe v23.0 analysis. This converted data was searched against cell surface proteome atlas (CSPA)[70] filtered UniProtKB mouse proteome (1296 entries with 50% decoy sequence incorporated). A modified LFQ-MBR workflow was used in MSfragger to identify biotinylated surface proteins only. A “strict-trypsin” enzyme specificity with 20 ppm tolerance is used for identification. The variable modification used were methionine oxidation (+15.9949, maximum 3 occurrences) and NHS-LC-Biotinylation of Lysine (+339.1617, maximum 1 occurrence). The cysteine carbamidomethylation (+57.02146) was selected as fixed modification with 1% false discovery rate cutoff for PSM, peptides, and proteins. MS1 based quantification of surface biotinylated proteins were performed with match between run (MBR) algorithm enabled. The extracted intensities were curated and utilized in MetaboAnalyst v6.0 for differential analysis

#### Data analysis and statistical methods

The normalised intensities of the protein groups quantified in the samples were extracted from Spectronaut. The processed data were visualised via principal component analysis (PCA), and a clustering heatmap was generated using the pheatmap package in R. The statistical evaluation of the identified proteins was done using a two-tailed, unpaired Student’s t-test employing multiple testing correction. The candidates were filtered out by adjusted p-value (<0.05) and absolute log2 fold change of 0.58 and were visualised using a volcano plot. The unique and overlapping DEPs identified in the muscle and brain samples were visualised using an Upset plot generated using the UpSetR package (Conway et al., 2017) from the Comprehensive R Archive Network (CRAN). Over-representation analysis (ORA) of the DEPs was performed in the ShinyGO (v0.85) online tool using Reactome as the gene ontology database. The enriched pathways were represented using a chord plot.

For the MRM analysis, firstly, the transitions with high missing values (in 7 or more samples) were removed; the remaining transitions were statistically evaluated using an unpaired two-sided Student’s t test with 5 % permutation-based FDR in Perseus (v2.0.7.0).

### 7. Immunohistochemistry

Post-harvesting, half of the mouse brain was fixed in 4% paraformaldehyde solution (PFA) for 24 h, followed by dehydration in 30% sucrose. Tissues were fixed in a cryo-matrix (Polysciences, 19636-1) and sectioned at 30 μm using a cryomicrotome (SLEE MNT). The sections were washed 3 times with 1X PBS. The sections were incubated with 3% H_2_O_2_ for 5 min and later washed 3 times with 1X PBS. The sections were blocked with a blocking buffer containing 1% BSA, 10% goat serum, and 0.1% Triton X-100 in 1X PBS for 1 h at room temperature. The primary antibody was prepared in blocking buffer and incubated with the sections at 4°C overnight. The sections were rinsed 3 times with PBST (PBS-0.1% Tween-20) and incubated with an HRP-conjugated secondary antibody at room temperature for 1 h. The detection was performed using SignalStain® DAB Substrate Kit (CST, 8059) according to the manufacturer’s protocol. The sections were mounted onto the charged slides and mounted using DPX mounting reagent (SRL, 88147).

### 8. Immunofluorescence

Upon harvesting, the muscle tissues were embedded in cryomatrix media (Polysciences, 19636-1) and flash-frozen in cooled 2-methylbutane (Sigma, M32631). The tissue sections at 10 μm were obtained using a cryomicrotome (SLEE MNT) and collected on the charged glass slides (Avantor; 631-0108). The sections were fixed in 4% PFA and later blocked in 10% goat serum, 1% Tergitol (NP-40) in PBS for 1 h. The sections were incubated with the primary antibody dilutions in 10% goat serum and PBS at 4°C overnight. The slides were rinsed with PBST and incubated with fluorophore-conjugated secondary antibodies at room temperature for 1 h. The slides were later washed with PBS and stained with DAPI (ThermoFisher Scientific; H3570) for 10 min at room temperature. The slides were mounted using Fluoromount-G (Southern Biotech; 0100-01). The sections were imaged using an Olympus fluorescence microscope (Olympus BX63F), with a Hamamatsu ORCA-Flash 4.0 camera or Leica SP8 confocal microscope. The muscle fiber area was quantified using SMASH (Semi-automatic muscle analysis using segmentation of histology software), and the cross-sectional area was measured using the CellSens Dimension 1.16 software. To visualise NMJs, adjacent 30 µm cryosections were fixed in 2% PFA at 4°C for 2 h, rinsed with PBS, and blocked with 5% goat serum for 1 h at room temperature. Sections were then incubated overnight at 4°C with α-bungarotoxin in PBS to label postsynaptic acetylcholine receptors, followed by three PBS washes of 5 minutes each. Subsequently, sections were incubated overnight at 4°C with anti-synaptophysin primary antibody to label presynaptic terminals, followed by an Alexa Fluor 488-conjugated secondary antibody for 2 hours at room temperature. Sections were rinsed in PBS, counterstained with DAPI, post-fixed in 4% PFA, and mounted with Fluoromount-G.

For immunostaining, N2A and C2C12 cells were cultured on glass coverslips and, upon treatment, fixed with 4% PFA for 5 min at room temperature. The cells were blocked with 3% BSA and 0.1% Tween 20 in PBS for 1 h at room temperature. The cells were incubated with primary antibody dilutions prepared in blocking buffer and incubated for 2 h at room temperature. The coverslips were rinsed with PBS and incubated with fluorophore-conjugated secondary antibodies at room temperature for 1 h. The coverslips were later washed with PBS and stained with DAPI for 10 min at room temperature. The coverslips were rinsed with MilliQ water, air-dried, and later mounted using Fluoromount-G. The coverslips were imaged on a Leica SP8 confocal microscope using a 63X objective. Image processing was done using Leica LASX software. Fluorescence intensity measurements were performed using Fiji.

### 9. ɑSynuclein purification, labelling and fibril preparation

The ɑSyn protein was expressed using the pT7-7 plasmid (Addgene, 36046) and transformed into BL21 (DE3) *E. coli cells.* Proteins were purified using the protocol as previously reported[125] and briefly described here. BL21 (DE3) *E. coli* cells were grown in LB medium at 37◦C and 180 rpm, and induction with 1 mM IPTG was performed at an OD_600_ of 0.6, with incubation for 4 h for protein expression. The cells were collected after centrifugation at 6000×g for 30 min, and the pellet was resuspended in lysis buffer, 50 mM Tris, 10 mM EDTA, and 150 mM NaCl, pH 8. To the resuspended cells, 1 mM PMSF and 0.5 mg/mL lysozyme were added, and the mixture was incubated for 30 min. Further, the suspension was sonicated at 50 % amplitude with a 10 min cycle of 45s ‘on’ and 15s ‘off’, and the lysate obtained was heated at 95◦C for 20 min. After centrifugation at 14000×g for 30 min, the supernatant was collected, and streptomycin sulfate was added at a concentration of 10 mg/mL and incubated at 4◦C for 30 min with occasional shaking. The solution was then centrifuged at 14000×g for 30 min, and the supernatant was collected. The protein precipitation was performed with ammonium sulfate at 360 mg/ml, the solution was incubated for 1 h at 4◦C and later centrifuged at 14000×g for 30min. The pellet was collected, resuspended in a (1:1) mixture of 100 mM ammonium acetate and 100 % ethanol. The mixture was incubated on ice for 10 min, then centrifuged at 14000×g for 10 min. This step was repeated thrice. The obtained pellet was finally resuspended in 100 mM ammonium acetate, flash-frozen in liquid nitrogen, and lyophilized. The lyophilized αSyn protein was further processed following the previously published protocol[126]. The protein was resuspended in 20 mM glycine NaOH buffer, pH 7.4. To dissolve the protein, a few drops of 0.2 M NaOH were added, and the solution was incubated until it became clear. Later, the solution pH was adjusted to 7.4 with 2 M HCl. To remove any high-order aggregates, the solution was passed through a 100 kDa cutoff filter (Millipore, UFC9100). The flow-through, consisting majorly of αSyn monomeric protein, was collected. The solution was buffer-exchanged into 20 mM sodium phosphate buffer, pH 7.4, using a 3 kDa cutoff filter (Millipore, UFC9010). The protein concentration was measured using Pierce™ BCA Protein Assay Kit (Thermo, A55864).

#### Protein Labelling

ɑSynuclein was labelled using conjugation with NHS-Rhodamine (Thermo 46406). The NHS-Rhodamine was dissolved at 10 mg/ml in DMSO. To label 200 μM of αSyn, 10 molar excess NHS-Rhodamine was added in 20 mM Sodium Phosphate buffer, pH 7.4, and incubated in the dark for 2 h at 4°C. The non-reacted NHS-Rhodamine was removed by passing the mixture through a desalting column (BIO-RAD, 7322010). The protein concentration was estimated with the Pierce™ BCA Protein Assay Reagent Kit (Thermo, A55864). The labelled protein aliquots were prepared, flash-frozen, and stored at -80°C until use.

#### Fibrillation

To prepare ɑSynuclein fibrils, 500 μL of 300 μM αSyn protein, diluted in 20 mM sodium phosphate buffer, pH 7.4, 0.1% sodium azide, along with a 3 mm glass bead, was added to a 2 mL micro centrifuge tube. The tube was incubated in an orbital shaker at 200 rpm, 37°C for 7 days to induce αSyn aggregation. To prepare fluorescently labelled αSyn fibrils, 0.5 % NHS-Rhodamine-labelled αSyn protein was added to the above mixture and incubated as described above. The fibrillation was confirmed by Transmission Electron Microscopy and Atomic Force Microscopy. To prepare preformed fibrils (PFFs), the αSyn fibrils were probe-sonicated for 90 sec at 20 % amplitude in a 0.5 sec on/off cycle. The PFFs were prepared and confirmed using Dynamic Light Scattering (DLS) and Atomic Force Microscopy (AFM).

### 10. Biolayer Interferometry

To determine αSyn interactions with TFRC, Biolayer Interferometry (BLI) (Octet RED96e) was performed using a previous protocol [20]. Initially, 15 μg of recombinant human Transferrin Receptor (TFRC) (Cat: 2474-TR-050, R&D Biosystems) was immobilized to AR2G (Amine Reactive Second-Generation) (Cat: 18–5092, Sartorius) sensors through amine coupling. The AR2G sensors were hydrated in Milli-Q water for 10 min. The sensor was activated with 20 mM EDC (1-ethyl-3-(3-(dimethylamino)propyl) carbodiimide) and 10 mM NHS (N-hydroxysuccinimide) in Milli-Q water for 300 s. TFRC protein was reconstituted in 10 mM sodium acetate buffer, pH 5, and loading was performed until saturation was reached. Quenching was performed by dipping the sensor into 1 M ethanolamine for 300 s. The baseline, association, and dissociation reactions were carried out in 20 mM sodium phosphate with 150 mM NaCl pH 7.4 for 120 s each at 25 ◦C, with constant agitation at 1000 rpm. The regeneration and neutralization reactions were conducted for 5 s each and repeated thrice in 50 mM NaOH and sodium phosphate buffer, respectively. The reactions were conducted in black polypropylene 96-well microplates. Data was analyzed using Data Analysis (FORTÉBIO) software with Savitzky−Golay filtering. The binding curves were reference-corrected, and different fitting models were tested, with the best fit observed in the 2:1 ligand-binding model for αSyn and transferrin.

### 11. Cell Culture

N2A and C2C12 cells were cultured in DMEM media supplemented with 10 % fetal bovine serum and 1X antibiotic cocktail at 37°C under 5% CO2. C2C12 myoblasts were differentiated to myotubes in differentiation media with 2% horse serum supplemented in DMEM media. Initially, cells were cultured in DMEM complete media, and after 24h, differentiation media were added. The cells were cultured in differentiation media until myotube formation was observed. N2A cells were differentiated using differentiation media with 10 μM retinoic acid and 1% fetal bovine serum supplemented in DMEM media. Differentiation media was added after 24h of maintenance in DMEM complete media.

#### Bodipy-C11 Lipid Peroxidation Assay

To quantify lipid peroxidation in N2A and C2C12 cells lipid peroxidation assay was performed. For the experiment, cells were seeded at 50,000 cells per well in 96-well plates in complete DMEM and allowed to adhere overnight. The next day, cells were washed with 1X PBS and incubated with 5 µM BODIPY™ 581/591 C11(Invitrogen, D3861) in serum free media for 30 min at 37°C. Following incubation, cells were washed twice with 1XPBS to remove excess probe and subsequently treated with 5 μM Erastin, 2.5 μM αSyn pre-formed fibrils, and 10 μM ferrostatin-1, and incubated for 24h at 37°C. Upon incubation, fluorescence intensities for both the oxidized (ex: 485 nm, em: 520 nm) and reduced (ex: 581 nm, em: 591 nm) dye were measured using SpectraMax i3x Multi-Mode Microplate Reader (Molecular Devices). Lipid peroxidation was quantified as the ratiometric shift in green-to-red fluorescence intensity, with an increase in the ratio indicating oxidation of polyunsaturated fatty acids. Data were normalized to vehicle-treated control groups, and background fluorescence was subtracted using unstained cell controls.

#### FerroOrange Assay

To measure intracellular ferrous iron, the FerroOrange (Cell Signaling Technology, 36104) assay was performed according to the manufacturer’s instructions. Briefly, cells were seeded onto glass coverslips in a 12-well plate in complete DMEM at 37°C. Cells were treated with 2.5 μM αSyn PFFs and 1X PBS as a vehicle control, and incubated for 24h at 37°C. After incubation, the cells were washed twice with 1X PBS, and a 1 µmol/L solution of FerroOrange in serum-free media was added and incubated for 30 min at 37°C. Nuclei were counterstained with DAPI. Cells were examined under a confocal microscope (Leica SP8) and images were captured. FerroOrange fluorescence intensity was measured using LASX software.

### 12. Statistical Analysis

All the experiments are performed at least in triplicate. Data were analyzed using GraphPad Prism 10. Data are shown as mean ± SEM, with at least 3 biological replicates per experimental condition. Statistical analysis employed a parametric, two-tailed, unpaired t-test or ANOVA followed by Tukey’s multiple comparison test with *P < 0.05, **P < 0.01, ***P < 0.001, ****P < 0.0001.

## Data Availability

All data corresponding to the manuscript are available in the manuscript or as supplementary information.

## Disclosures

Authors declare no competing interests.

## Authors Contributions

KSB: Conceptualization, methodology, investigation, visualization, data curation, validation, formal analysis, and writing (original draft)

JS: Investigation, methodology, data curation, formal analysis, and validation.

NK: Investigation, methodology, data curation, formal analysis.

AB: Investigation, methodology, data curation, formal analysis.

SS: Investigation, methodology.

SJM: Conceptualization, resources, supervision

TKM: Conceptualization, investigation, methodology, funding acquisition, resources, supervision, project administration, visualization, and writing (original draft, review, and editing).

All the authors read and approved the manuscript.

## Funding and additional information

KSB and JS thank the INSPIRE fellowship from DST (Department of Science and Technology, Government of India), Naman thanks the senior research fellowship (SRF) from CSIR (Council of Scientific and Industrial Research), Ankit thanks the SRF fellowship from DBT (Department of Biotechnology), Sandhini Saha thank Indian Council for Medical Research (ICMR) for the SRF fellowship, Tushar K Maiti and Sam J Mathew thank the Regional Centre for Biotechnology (RCB) for funding ( No. RCB/C0100)

## Acknowledgements

Authors acknowledge the small animal facility (SAF) for animal work. We thank Dr. Hitesh Kamboj (RCB) for help with animal studies. Authors thank Regional Centre Biotechnology (RCB) for state-of-the-art central instrumentation facility (CIF) and mass spectrometry facility. Advanced Technology Platform Centre (ATPC) at RCB for BLI facility. Authors acknowledge the support of DBT e-Library Consortium (DeLCON) and One Nation One Subscription (ONOS) for providing access to e-resources.

## Notes

### Competing Interest Statement

The authors have declared no competing interest.

