## Supplementary document for "Ferroptosis links α-synuclein pathology across brain and skeletal muscle in Parkinson’s disease"

***Running Title: α-Synuclein–TFRC Ferroptosis Axis in Parkinson's Disease***

#### Supplementary Materials

### 1. Supplementary Figures

#### Supplementary Figure 1

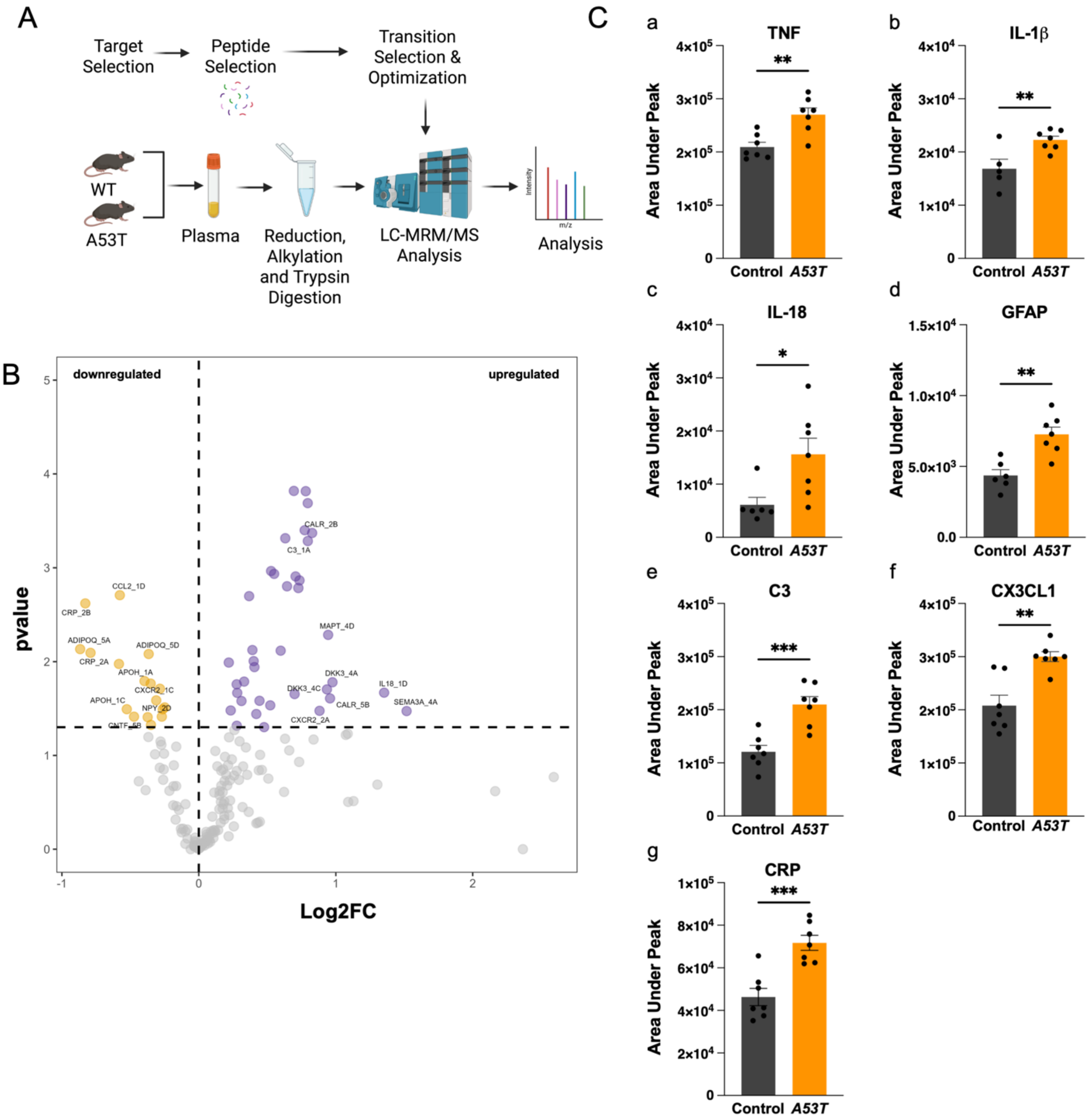

**Figure S1. Plasma multiple reaction monitoring (MRM) analysis of A53T mice reveals altered inflammatory markers.**

A. Schematic workflow of MRM utilizing plasma samples collected from both control and A53T mice.

B. Volcano plot depicting comparison between identified plasma proteins in control vs A53T mice with log2 fold change (FC) in the x-axis and $-\log_{10}$ p-value in y-axis, C. Quantification of plasma protein, namely (a) TNF, (b) IL-1β, (c) IL-18, (d) GFAP, (e) C3, (f) CX3CL1, and (g) CRP, represented as the area under the peak in control and A53T mice.

Data presented as mean ± SEM; an unpaired two-tailed Student's t-test was employed, with $***P < 0.001$, $**P < 0.01$, and $*P < 0.05$.

**Supplementary Figure 2**

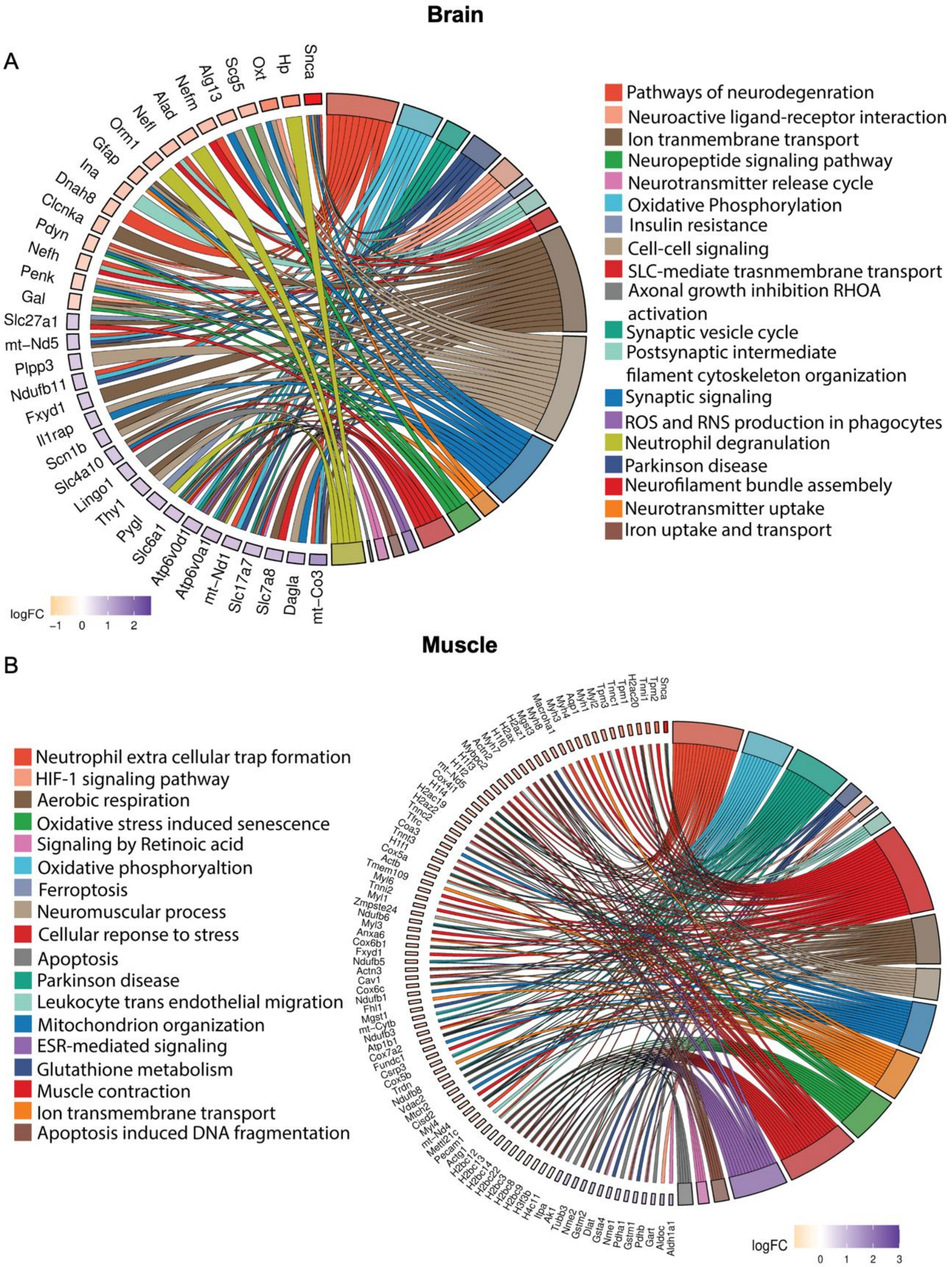
Brain
A
Snca
Hp
Oxt
Scg5
Alg13
Nefm
Alad
Nefl
Orm1
Gfap
Ina
Dnah8
Clcnka
Pdyn
Nefh
Penk
Gal
Slc27a1
mt−Nd5
Plpp3
Ndufb11
Fxyd1
Il1rap
Scn1b
Slc4a10
Lingo1
Thy1
Pygl
Slc6a1
Atp6v0d1
Atp6v0a1
mt−Nd1
Slc17a7
Slc7a8
Dagla
mt−Co3
logFC
−1 0 1 2
Pathways of neurodegenration
Neuroactive ligand-receptor interaction
Ion tranmembrane transport
Neuropeptide signaling pathway
Neurotransmitter release cycle
Oxidative Phosphorylation
Insulin resistance
Cell-cell signaling
SLC-mediate trasnmembrane transport
Axonal growth inhibition RHOA activation
Synaptic vesicle cycle
Postsynaptic intermediate filament cytoskeleton organization
Synaptic signaling
ROS and RNS production in phagocytes
Neutrophil degranulation
Parkinson disease
Neurofilament bundle assembely
Neurotransmitter uptake
Iron uptake and transport
Muscle
B
Neutrophil extra cellular trap formation
HIF-1 signaling pathway
Aerobic respiration
Oxidative stress induced senescence
Signaling by Retinoic acid
Oxidative phosphoryaltion
Ferroptosis
Neuromuscular process
Cellular reponse to stress
Apoptosis
Parkinson disease
Leukocyte trans endothelial migration
Mitochondrion organization
ESR-mediated signaling
Glutathione metabolism
Muscle contraction
Ion transmembrane transport
Apoptosis induced DNA fragmentation
logFC
0 1 2 3

**Figure S2. Pathway enrichment of Differentially Expressed Proteins.**

A, B. Chord plot representing GO (gene ontology) biological pathway enrichment from differentially expressed proteins (DEPs) in the brain (A) and gastrocnemius muscle (B) of A53T mice compared to control.

**Supplementary Figure 3**

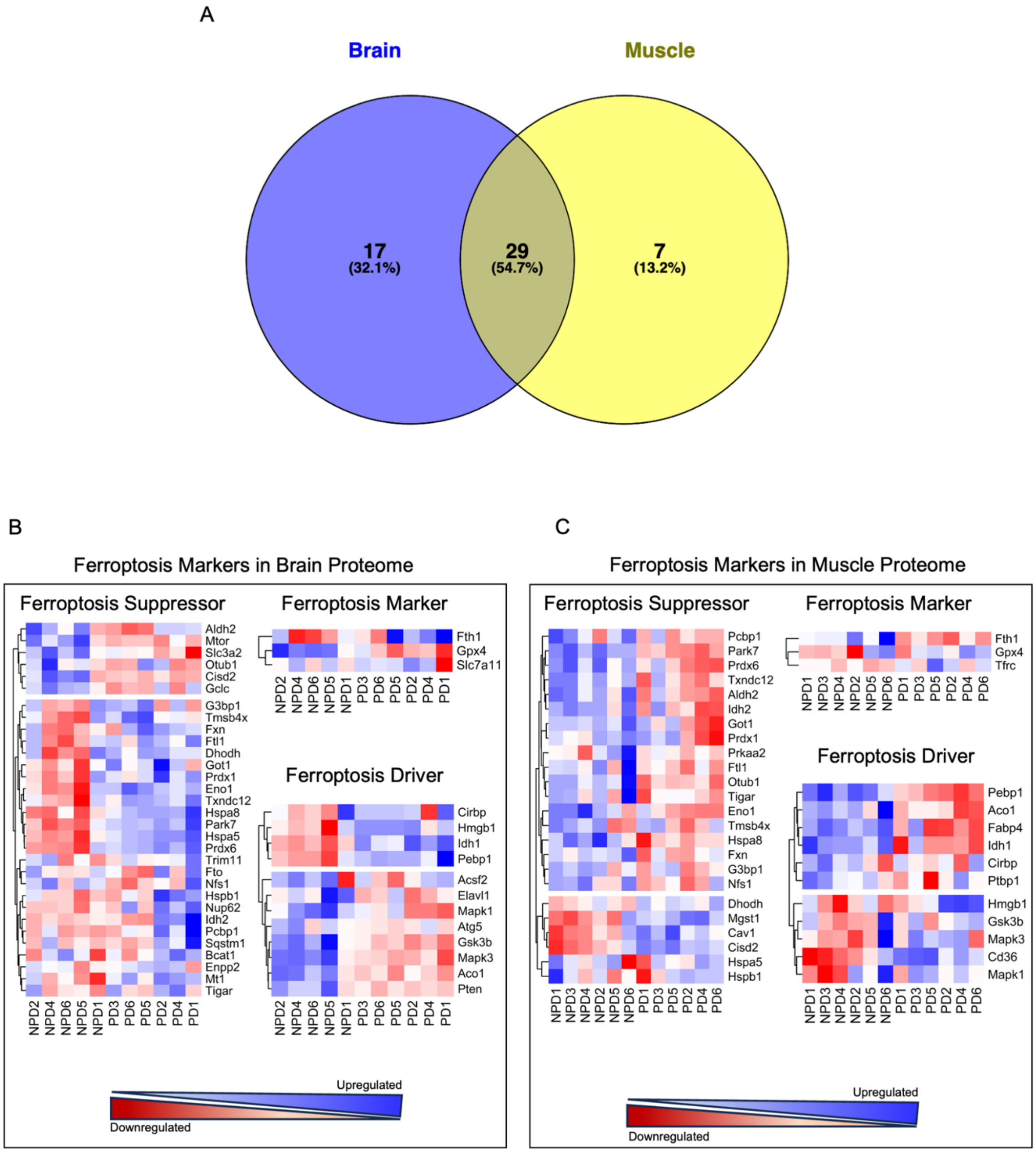


**Figure S3. Proteomic profiling reveals ferroptosis-associated signatures in the brain and the skeletal muscle.**

A. Venn diagram representing common and unique ferroptosis-associated proteins identified in the brain and muscle SWATH-MS data.

B, C. Ferroptosis-associated proteins identified in the brain (B) and gastrocnemius muscle (C) are categorized into three functional groups: drivers, suppressors, and markers using the ferroptosis proteins database. The intensity distribution of segregated proteins is represented as a heatmap.

**Supplementary Figure 4**

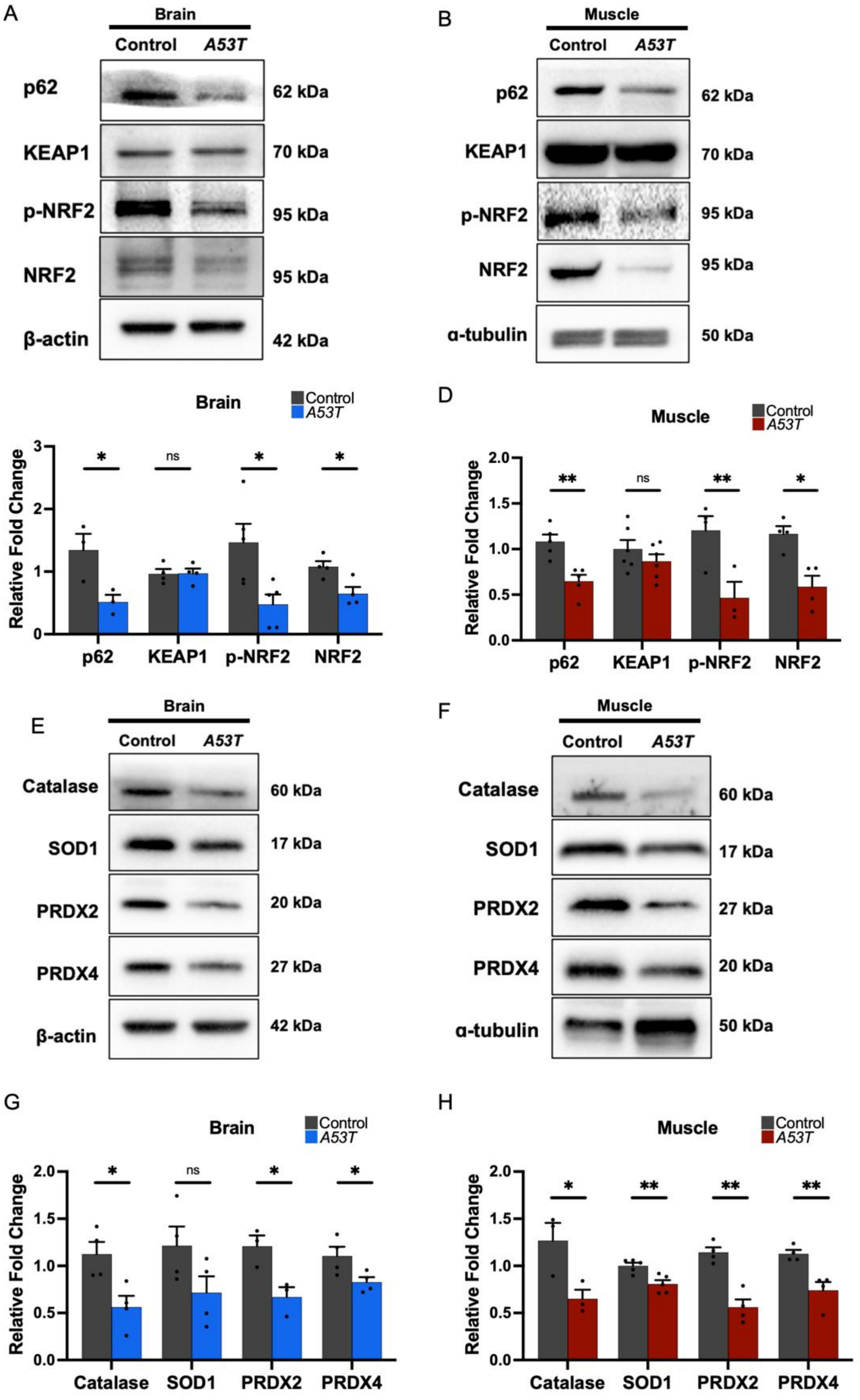
A
Brain
Control
A53T
p62
KEAP1
p-NRF2
NRF2
β-actin
62 kDa
70 kDa
95 kDa
95 kDa
42 kDa
B
Muscle
Control
A53T
p62
KEAP1
p-NRF2
NRF2
α-tubulin
62 kDa
70 kDa
95 kDa
95 kDa
50 kDa
C
Brain
Control
A53T
Relative Fold Change
0
1
2
3
*
ns
*
*
p62
KEAP1
p-NRF2
NRF2
D
Muscle
Control
A53T
Relative Fold Change
0.0
0.5
1.0
1.5
2.0
**
ns
**
*
p62
KEAP1
p-NRF2
NRF2
E
Brain
Control
A53T
Catalase
SOD1
PRDX2
PRDX4
β-actin
60 kDa
17 kDa
20 kDa
27 kDa
42 kDa
F
Muscle
Control
A53T
Catalase
SOD1
PRDX2
PRDX4
α-tubulin
60 kDa
17 kDa
27 kDa
20 kDa
50 kDa
G
Brain
Control
A53T
Relative Fold Change
0.0
0.5
1.0
1.5
2.0
*
ns
*
*
Catalase
SOD1
PRDX2
PRDX4
H
Muscle
Control
A53T
Relative Fold Change
0.0
0.5
1.0
1.5
2.0
*
**
**
**
Catalase
SOD1
PRDX2
PRDX4

**Supplementary Figure 4. A53T mice exhibit suppression of NRF2 antioxidant signaling**

A-D. Representative western blot analysis for p62, KEAP1, p-NRF2, NRF2, beta-actin, and alpha tubulin (loading control) using protein lysates from the brain (A) and gastrocnemius muscle (B), their densitometric quantification (C) and (D), respectively.

E-H. Representative western blot analysis for Catalase, SOD1, PRDX2, PRDX4, beta-actin, and alpha tubulin (loading control) using protein lysates from the brain (E) and gastrocnemius muscle (F), their densitometric quantification (G) and (H), respectively.

Data presented as mean ± SEM; an unpaired two-tailed Student's t-test was employed, with $**P < 0.01$, and $*P < 0.05$.

**Supplementary Figure 5**

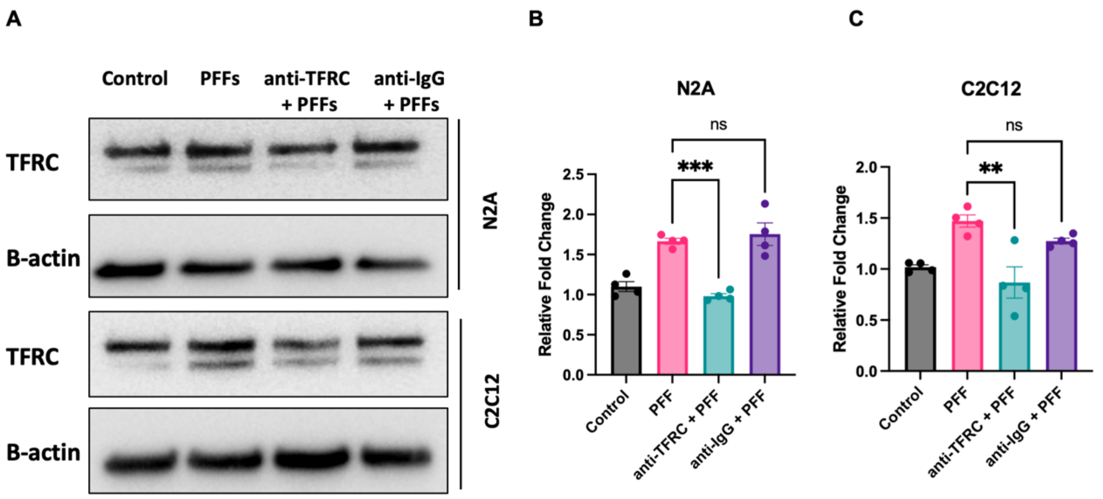


**Supplementary Figure 5. α-Synuclein induces TFRC expression.**

A. Representative western blot analysis for TFRC and beta actin (loading control) using protein lysates from N2A and C2C12 cells upon αSyn-PFF treatment (A) and densitometric analysis for N2A (B) and C2C12 cells (C). (n = 4 independent experiments).

Data is shown as mean ± sem, n = 4 independent experiments. One-way ANOVA followed by Tukey's correction, ***$P < 0.001$ and **$P < 0.01$.

#### 2. Supplementary Tables

##### Table S1: List of antibodies

| Reagents | Source | Cat. Number |
|---|---|---|
| **Primary Antibodies** | | |
| α-synuclein | Thermo Fisher Scientific | PA5-85791 |
| pS129 α-synuclein | Cell Signaling Technology | 23706S |
| Tyrosine Hydroxylase | Abclonal | A0028 |
| NCAM | Abclonal | A11770 |
| MuSK | Abclonal | A2591 |
| AChR | Abclonal | A10057 |
| LRP4 | Abclonal | A25225 |
| TFRC | Cell Signaling Technology | 55487 |
| FTH | Abclonal | A19544 |
| DMT1 | Abclonal | A10231 |
| SLC7A11 | Abclonal | A13685 |
| GPX4 | Abclonal | A1933 |
| GPX1 | Abclonal | A1110 |
| NRF2 | Proteintech | 98020000 |
| p-NRF2 | Abclonal | AP1133 |
| KEAP1 | Abclonal | A1820 |
| p62 | Cell Signaling Technology | 7695 |
| 4-HNE | Abclonal | A26085 |
| Catalase | Abclonal | A11220 |
| SOD1 | Abclonal | A0274 |
| Prdx1 | Abclonal | A13524 |
| Prdx4 | Abclonal | A18308 |
| Beta actin | Cloud Clone | PAB340Mi01 |
| Alpha tubulin | Cell Signaling Technology | 3873T |
| **Secondary Antibodies** | | |
| Alexa Flour 488 Goat anti-rabbit | Invitrogen | A11034 |
| Alexa Flour 594 Goat anti-mouse | Invitrogen | A11005 |
| Alexa Flour 594 phalloidin | Invitrogen | A12381 |
| Cy3 conjugated Goat anti-rabbit | Jackson ImmunoResearch Laboratories | 111-165-144 |
| Biotin conjugated Goat-anti-mouse | Jackson ImmunoResearch Laboratories | 115-065-020 |

| | | |
|---|---|---|
| Biotin conjugated Goat anti- rabbit | Jackson ImmunoResearch Laboratories | 111-065-144 |
| Cy2 conjugated streptavidin | Jackson ImmunoResearch Laboratories | 016-220-084 |
| Goat anti-rabbit, HRP | Thermo Fisher Scientific | 31460 |
| Goat anti-mouse, HRP | Thermo Fisher Scientific | 31430 |

**Table S2: Differentially expressed proteins between control and A53T mice in the brain.**

| S. No. | UniProt IDs | Genes | Q value | Fold Change |
|---|---|---|---|---|
| 1 | P22005 | Penk | 1.72E-34 | 1.524067 |
| 2 | O55042 | Snca | 1.36E-27 | 6.268199 |
| 3 | P12961 | Scg5 | 2.90E-27 | 1.867391 |
| 4 | P10518 | Alad | 4.33E-26 | 1.785965 |
| 5 | Q5FWK3 | Arhgap1 | 6.01E-23 | 0.64089 |
| 6 | P08553 | Nefm | 2.43E-21 | 1.799772 |
| 7 | Q3TXX4 | Slc17a7 | 3.46E-21 | 0.60681 |
| 8 | P08551 | Nefl | 3.51E-21 | 1.758826 |
| 9 | P46660 | Ina | 6.08E-20 | 1.695094 |
| 10 | Q5DTL9 | Slc4a10 | 3.38E-19 | 0.65082 |
| 11 | Q8BI08 | Mal2 | 7.44E-18 | 0.505655 |
| 12 | Q9CPZ8 | Cmc1 | 2.62E-17 | 1.653871 |
| 13 | Q9CPU4 | Mgst3 | 6.93E-17 | 0.627037 |
| 14 | P03995 | Gfap | 1.12E-16 | 1.728764 |
| 15 | P55088 | Aqp4 | 4.76E-16 | 0.602456 |
| 16 | Q99JY8 | Plpp3 | 1.15E-15 | 0.658542 |
| 17 | P03888 | Mtnd1 | 2.43E-15 | 0.613658 |
| 18 | P31648 | Slc6a1 | 2.86E-15 | 0.641369 |
| 19 | P51863 | Atp6v0d1 | 7.71E-15 | 0.619885 |
| 20 | Q8VDG6 | Map3k21 | 1.11E-14 | 1.58019 |
| 21 | P19246 | Nefh | 1.23E-14 | 1.54227 |
| 22 | Q9Z1G4 | Atp6v0a1 | 1.49E-14 | 0.616236 |
| 23 | Q8BGN3 | Enpp6 | 2.27E-14 | 0.50621 |
| 24 | Q60714 | Slc27a1 | 2.31E-14 | 0.660926 |
| 25 | P01831 | Thy1 | 3.39E-14 | 0.643049 |
| 26 | P00416 | mt-Co3 | 1.33E-13 | 0.436636 |
| 27 | P01868;P01869 | Ighg1 | 2.18E-13 | 2.553118 |
| 28 | Q8BRH4 | Kmt2c | 8.63E-13 | 1.516969 |
| 29 | Q9ESJ4 | Nckipsd | 8.74E-13 | 0.570307 |
| 30 | P97952 | Scn1b | 3.43E-12 | 0.651733 |
| 31 | Q9D8C3 | Alg13 | 3.93E-12 | 1.813981 |

| | | | | |
|---|---|---|---|---|
| 32 | O09111 | Ndufb11 | 8.10E-12 | 0.657125 |
| 33 | P03921 | Mtnd5 | 8.31E-12 | 0.659508 |
| 34 | Q9DCT1 | Akr1e2 | 1.93E-11 | 0.518564 |
| 35 | P47212 | Gal | 2.33E-11 | 1.510131 |
| 36 | Q61361 | Bcan | 1.47E-10 | 0.384124 |
| 37 | Q9D1X9 | Tmem50b | 3.65E-10 | 0.624192 |
| 38 | Q9Z239 | Fxyd1 | 4.10E-10 | 0.656166 |
| 39 | Q0VBD2 | Mcm10 | 5.28E-10 | 0.592314 |
| 40 | Q9QXW9 | Slc7a8 | 1.06E-09 | 0.584702 |
| 41 | D3YZF7 | Vsig10l | 1.76E-09 | 1.871342 |
| 42 | Q9D114 | Hddc3 | 2.39E-09 | 1.516867 |
| 43 | G5E870 | Trip12 | 4.92E-09 | 0.503063 |
| 44 | Q9Z306 | Slc22a4 | 8.14E-09 | 0.668753 |
| 45 | Q9D1T0 | Lingo1 | 1.24E-08 | 0.650337 |
| 46 | O35417 | Pdyn | 3.20E-08 | 1.562481 |
| 47 | Q9CXX9 | Cuedc2 | 1.23E-07 | 2.946908 |
| 48 | P01872 | Ighm | 1.40E-07 | 0.518499 |
| 49 | Q61730 | Il1rap | 6.58E-07 | 0.655032 |
| 50 | Q14C51 | Ptcd3 | 1.57E-06 | 0.667071 |
| 51 | G5E869 | Znf142 | 3.35E-06 | 0.648947 |
| 52 | Q61646 | Hp | 4.06E-06 | 2.903024 |
| 53 | B1AXV0 | Frrs1l | 1.08E-05 | 0.525261 |
| 54 | P35454 | Oxt | 1.42E-05 | 2.870132 |
| 55 | Q8BRF7 | Scfd1 | 1.44E-05 | 0.549162 |
| 56 | B2RQE8 | Arhgap42 | 1.64E-05 | 0.432304 |
| 57 | Q91X72 | Hpx | 9.10E-05 | 1.508673 |
| 58 | P02089 | Hbb-b2 | 9.47E-05 | 0.586529 |
| 59 | Q9WUB7 | Clcnka | 0.000135 | 1.631736 |
| 60 | Q9ET01 | Pygl | 0.000229 | 0.642613 |
| 61 | Q64288 | Omp | 0.000237 | 0.477811 |
| 62 | A3KGS3 | Ralgapa2 | 0.000297 | 0.12558 |
| 63 | Q60590 | Orm1 | 0.0003 | 1.75181 |
| 64 | Q91VX2 | Ubap2 | 0.001468 | 1.56557 |
| 65 | O08692 | Ngp | 0.003697 | 1.965417 |
| 66 | Q91XQ0 | Dnah8 | 0.008762 | 1.65908 |
| 67 | Q9D777 | Tnfsf13 | 0.014854 | 0.118764 |
| 68 | Q6WQJ1 | Dagla | 0.018777 | 0.568786 |
| 69 | E9Q7M2 | Tsc22d2 | 0.019334 | 1.538858 |
| 70 | Q9DB43 | Zfpl1 | 0.019718 | 0.636122 |
| 71 | Q9DB75 | Cdip1 | 0.030501 | 0.646137 |

**Table S3: Differentially expressed proteins between control and A53T mice in the skeletal muscle.**

| S. No. | UniProt IDs | Genes | Q value | Fold Change |
|---|---|---|---|---|
| 1 | Q61941 | Nnt | 1.21E-37 | 2.287381 |
| 2 | Q9CYT6 | Cap2 | 1.66E-36 | 2.0137 |
| 3 | Q99JB8 | Pacsin3 | 1.52E-35 | 1.88324 |
| 4 | P62806 | H4c1 | 8.10E-35 | 2.775705 |
| 5 | Q64523 | H2ac20 | 3.75E-34 | 2.73928 |
| 6 | P19783 | Cox4i1 | 5.10E-33 | 1.801929 |
| 7 | Q9D051 | Pdhb | 1.07E-32 | 0.607478 |
| 8 | P45591 | Cfl2 | 2.60E-30 | 1.554683 |
| 9 | Q9CPU4 | Mgst3 | 4.84E-30 | 1.993297 |
| 10 | Q91ZJ5 | Ugp2 | 7.71E-30 | 0.65992 |
| 11 | Q01768 | Nme2 | 8.05E-30 | 0.638039 |
| 12 | P02301 | H3-5 | 9.14E-30 | 2.635689 |
| 13 | P03921 | Mtnd5 | 1.57E-29 | 1.802324 |
| 14 | P0C0S6 | H2az1 | 4.09E-29 | 1.986787 |
| 15 | P40124 | Cap1 | 5.33E-29 | 2.244426 |
| 16 | O54962 | Banf1 | 9.32E-29 | 1.876678 |
| 17 | Q3UIU2 | Ndufb6 | 2.10E-27 | 1.63151 |
| 18 | P15626 | Gstm2 | 4.30E-27 | 0.636948 |
| 19 | A2AUC9 | Klhl41 | 4.57E-27 | 1.585802 |
| 20 | Q9CPQ1 | Cox6c | 6.07E-27 | 1.588014 |
| 21 | P19123 | Tnnc1 | 1.25E-26 | 2.646011 |
| 22 | O55042 | Snca | 2.36E-26 | 12.19922 |
| 23 | P12787 | Cox5a | 2.95E-26 | 1.68676 |
| 24 | Q9WUZ5 | Tnni1 | 7.93E-26 | 2.956924 |
| 25 | P0DN34 | Ndufb1 | 9.73E-26 | 1.585184 |
| 26 | P10853 | H2bc7 | 1.13E-25 | 1.998082 |
| 27 | Q5XKE0 | Mybpc2 | 1.69E-25 | 1.855169 |
| 28 | P51667 | Myl2 | 2.09E-25 | 2.2292 |
| 29 | Q9ERK4 | Cse1l | 2.09E-25 | 2.013483 |
| 30 | Q9CQH3 | Ndufb5 | 2.39E-25 | 1.600425 |
| 31 | Q9QZ47 | Tnnt3 | 2.39E-25 | 1.702561 |
| 32 | P43277 | H1-3 | 5.94E-25 | 1.813417 |
| 33 | Q9D6J5 | Ndufb8 | 6.41E-25 | 1.537863 |
| 34 | P35486 | Pdha1 | 6.73E-25 | 0.62144 |
| 35 | Q6NT99 | Dusp23 | 1.03E-24 | 0.64611 |
| 36 | Q9R0Y5 | Ak1 | 1.03E-24 | 0.646173 |
| 37 | Q60930 | Vdac2 | 1.29E-24 | 1.520649 |
| 38 | Q99LX0 | Park7 | 2.28E-24 | 0.666772 |

| | | | | |
|---|---|---|---|---|
| 39 | Q9QZQ8 | Macroh2a1 | 4.42E-24 | 2.074027 |
| 40 | P10922 | H1-0 | 1.03E-23 | 1.934592 |
| 41 | P97457 | Myl11 | 1.11E-23 | 1.678238 |
| 42 | P15864 | H1-2 | 1.24E-23 | 1.807274 |
| 43 | Q9D783 | Klhl40 | 6.44E-23 | 1.511184 |
| 44 | Q8BMF4 | Dlat | 7.65E-23 | 0.632167 |
| 45 | P08074 | Cbr2 | 9.92E-23 | 0.592604 |
| 46 | Q02013 | Aqp1 | 1.09E-22 | 2.122662 |
| 47 | E9Q9K5 | Trdn | 2.50E-22 | 1.564414 |
| 48 | P43274 | H1-4 | 2.89E-22 | 1.78116 |
| 49 | P48758 | Cbr1 | 3.78E-22 | 0.611918 |
| 50 | Q64737 | Gart | 1.39E-21 | 0.607358 |
| 51 | P05977 | Myl1 | 2.50E-21 | 1.645468 |
| 52 | P19536 | Cox5b | 2.76E-21 | 1.566394 |
| 53 | P09542 | Myl3 | 3.87E-21 | 1.620365 |
| 54 | P14094 | Atp1b1 | 3.92E-21 | 1.581035 |
| 55 | P48771 | Cox7a2 | 4.61E-21 | 1.579672 |
| 56 | Q08857 | Cd36 | 4.67E-21 | 1.638107 |
| 57 | Q99MQ4 | Aspn | 1.05E-20 | 1.663148 |
| 58 | Q9JI91 | Actn2 | 1.11E-20 | 1.890052 |
| 59 | P24472 | Gsta4 | 1.19E-20 | 0.630043 |
| 60 | Q9CQB5 | Cisd2 | 1.40E-20 | 1.506114 |
| 61 | P56391 | Cox6b1 | 1.66E-20 | 1.606066 |
| 62 | Q9CY50 | Ssr1 | 2.39E-20 | 1.536617 |
| 63 | P97447 | Fhl1 | 3.71E-20 | 1.584638 |
| 64 | Q9QZZ6 | Dpt | 5.26E-20 | 1.981552 |
| 65 | P32848 | Pvalb | 1.32E-19 | 0.667861 |
| 66 | Q64105 | Spr | 1.48E-19 | 0.625549 |
| 67 | P21614 | Gc | 1.63E-19 | 1.976399 |
| 68 | Q791V5 | Mtch2 | 1.74E-19 | 1.514889 |
| 69 | Q9CPZ8 | Cmc1 | 2.48E-19 | 1.594689 |
| 70 | O88990 | Actn3 | 2.71E-19 | 1.598455 |
| 71 | Q7TPR4 | Actn1 | 3.03E-19 | 1.525322 |
| 72 | P13541 | Myh3 | 3.69E-19 | 2.079695 |
| 73 | Q8K2Q5 | Chchd7 | 7.10E-19 | 1.561331 |
| 74 | Q9WV35 | Apobec2 | 8.68E-19 | 3.063208 |
| 75 | Q9CQZ6 | Ndufb3 | 9.84E-19 | 1.58114 |
| 76 | Q60605 | Myl6 | 1.04E-18 | 1.649693 |
| 77 | Q9CPU0 | Glo1 | 5.05E-18 | 0.51669 |
| 78 | P68134 | Acta1 | 6.15E-18 | 2.171091 |
| 79 | Q5SX40 | Myh1 | 9.96E-18 | 2.134033 |

| | | | | |
|---|---|---|---|---|
| 80 | P13542 | Myh8 | 3.19E-17 | 2.039396 |
| 81 | Q5SX39 | Myh4 | 3.38E-17 | 2.089582 |
| 82 | Q8VDQ1 | Ptgr2 | 7.65E-17 | 0.6624 |
| 83 | P21107 | Tpm3 | 8.14E-17 | 2.309641 |
| 84 | P53395 | Dbt | 2.30E-16 | 0.589843 |
| 85 | P00158 | Mt-Cyb | 2.50E-16 | 1.584525 |
| 86 | Q91Z83 | Myh7 | 2.56E-16 | 1.926062 |
| 87 | Q9Z0P5 | Twf2 | 3.22E-16 | 1.796717 |
| 88 | P01868 | Ighg1 | 3.50E-16 | 2.569945 |
| 89 | Q99J99 | Mpst | 5.09E-16 | 0.562728 |
| 90 | Q9CQS8 | Sec61b | 6.01E-16 | 1.515448 |
| 91 | Q8CHT0 | Aldh4a1 | 1.04E-15 | 0.371056 |
| 92 | P50462 | Csrp3 | 1.44E-15 | 1.576412 |
| 93 | Q80W54 | Zmpste24 | 3.32E-15 | 1.633253 |
| 94 | Q9DBS1 | Tmem43 | 3.82E-15 | 1.720631 |
| 95 | P58771 | Tpm1 | 5.74E-15 | 2.72644 |
| 96 | P24549 | Aldh1a1 | 5.82E-15 | 0.505763 |
| 97 | Q9D892 | Itpa | 5.84E-15 | 0.650788 |
| 98 | P59280 | Klhl8 | 5.97E-15 | 1.744982 |
| 99 | Q99MR8 | Mccc1 | 9.91E-15 | 0.523515 |
| 100 | Q9CZY3 | Ube2v1 | 1.00E-14 | 0.641906 |
| 101 | P49817 | Cav1 | 1.03E-14 | 1.590894 |
| 102 | P27661 | H2ax | 1.48E-14 | 1.976054 |
| 103 | P60710;P63260 | Actb;Actg1 | 8.63E-14 | 1.661186 |
| 104 | P01872 | Ighm | 8.91E-14 | 0.464196 |
| 105 | Q3UBX0 | Tmem109 | 1.42E-13 | 1.65022 |
| 106 | P14824 | Anxa6 | 4.83E-13 | 1.606315 |
| 107 | P27573 | Mpz | 1.06E-12 | 1.908767 |
| 108 | Q8C0L9 | Gpcpd1 | 1.43E-12 | 0.641272 |
| 109 | P09541 | Myl4 | 2.09E-12 | 1.503565 |
| 110 | Q9CR64 | Tmem167a | 2.23E-12 | 1.519263 |
| 111 | P58774 | Tpm2 | 2.34E-12 | 3.791595 |
| 112 | P35762 | Cd81 | 4.69E-12 | 1.740366 |
| 113 | P15532 | Nme1 | 7.55E-12 | 0.629363 |
| 114 | P20801 | Tnnc2 | 8.52E-12 | 1.756701 |
| 115 | Q8BH79 | Ano10 | 1.16E-11 | 1.597697 |
| 116 | Q9D2R6 | Coa3 | 1.55E-11 | 1.720897 |
| 117 | P03911 | Mtnd4 | 2.01E-11 | 1.501898 |
| 118 | Q9DBB8 | Dhdh | 2.22E-11 | 0.644508 |
| 119 | Q91VS7 | Mgst1 | 2.68E-11 | 1.584609 |
| 120 | Q8VCT4 | Ces1d | 3.71E-11 | 0.604344 |

| | | | | |
|---|---|---|---|---|
| 121 | Q3UV17 | Krt76 | 5.49E-11 | 0.644336 |
| 122 | Q8R1S0 | Coq6 | 9.46E-11 | 1.629231 |
| 123 | P43275 | H1-1 | 2.45E-10 | 1.700162 |
| 124 | P10649 | Gstm1 | 2.67E-10 | 0.620509 |
| 125 | P11087 | Col1a1 | 2.67E-10 | 1.578551 |
| 126 | Q9Z239 | Fxyd1 | 3.88E-10 | 1.600593 |
| 127 | Q62147 | Sspn | 5.44E-10 | 1.56332 |
| 128 | Q80UY1 | Carnmt1 | 1.40E-09 | 0.615633 |
| 129 | Q9ER42 | Barx1 | 2.87E-09 | 1.664201 |
| 130 | P01754 | Ighv1-62-3 | 6.43E-09 | 1.698096 |
| 131 | Q9WVL0 | Gstz1 | 1.57E-08 | 0.60292 |
| 132 | Q9DB70 | Fundc1 | 4.46E-08 | 1.576992 |
| 133 | Q8R2U4 | Ntmt1 | 4.84E-08 | 1.530569 |
| 134 | Q8CI94 | Pygb | 2.77E-07 | 0.636511 |
| 135 | P03987 | NaN | 3.22E-07 | 0.51044 |
| 136 | Q61838 | Pzp | 3.63E-07 | 0.599845 |
| 137 | P56695 | Wfs1 | 4.45E-07 | 2.256665 |
| 138 | P13412 | Tnni2 | 1.37E-06 | 1.647977 |
| 139 | P05063 | Aldoc | 4.28E-06 | 0.518053 |
| 140 | P97386 | Lig3 | 6.01E-06 | 0.320642 |
| 141 | Q61543 | Glg1 | 7.63E-06 | 1.584732 |
| 142 | Q8CFV9 | Rfk | 1.50E-05 | 0.612221 |
| 143 | Q3UVC0 | Ksr2 | 1.96E-05 | 0.252067 |
| 144 | P01867 | Ighg2b | 2.03E-05 | 1.775416 |
| 145 | Q61765 | Krt31 | 2.20E-05 | 0.460687 |
| 146 | P14426 | H2-D1 | 7.25E-05 | 0.568873 |
| 147 | Q61646 | Hp | 0.000111 | 1.918488 |
| 148 | Q9JLR1 | Sec61a2 | 0.000143 | 1.795323 |
| 149 | O35969 | Gamt | 0.000208 | 0.647235 |
| 150 | Q08481 | Pecam1 | 0.000273 | 1.495559 |
| 151 | Q9WV54 | Asah1 | 0.000338 | 0.632614 |
| 152 | P01636 | NaN | 0.001047 | 0.614635 |
| 153 | O35566 | Cd151 | 0.001801 | 1.751707 |
| 154 | Q497I4 | Krt35 | 0.002363 | 0.043573 |
| 155 | Q8BLU2 | Mettl21c | 0.002676 | 1.499939 |
| 156 | A6H6E2 | Mmrn2 | 0.003091 | 1.582507 |
| 157 | Q7TT45 | Rragd | 0.0037 | 0.642859 |
| 158 | Q3TEA8 | Hp1bp3 | 0.003824 | 1.495915 |
| 159 | Q80U28 | Madd | 0.006266 | 0.578651 |
| 160 | Q9R0Q3 | Tmed2 | 0.020825 | 1.894794 |
| 161 | Q62351 | Tfrc | 0.022028 | 1.731264 |

| | | | | |
|---|---|---|---|---|
| 162 | Q9ERD7 | Tubb3 | 0.023614 | 0.64195 |
| 163 | P01644 | NaN | 0.025683 | 1.506066 |
| 164 | Q8BZN4 | Nuak2 | 0.045487 | 0.63928 |

**Table S4. Ferroptosis-associated proteins identified in A53T vs Control mice in the brain.**

| S. No. | Genes | Gene Name | Group |
|---|---|---|---|
| 1 | Slc3a2 | 4F2_MOUSE | Ferr_Suppressor |
| 2 | Got1 | AATC_MOUSE | Ferr_Suppressor |
| 3 | Aldh2 | ALDH2_MOUSE | Ferr_Suppressor |
| 4 | Bcat1 | BCAT1_MOUSE | Ferr_Suppressor |
| 5 | Hspa5 | BIP_MOUSE | Ferr_Suppressor |
| 6 | Cisd2 | CISD2_MOUSE | Ferr_Suppressor |
| 7 | Eno1 | ENOA_MOUSE | Ferr_Suppressor |
| 8 | Enpp2 | ENPP2_MOUSE | Ferr_Suppressor |
| 9 | Fxn | FRDA_MOUSE | Ferr_Suppressor |
| 10 | Ftl1 | FRIL1_MOUSE | Ferr_Suppressor |
| 11 | Fto | FTO_MOUSE | Ferr_Suppressor |
| 12 | G3bp1 | G3BP1_MOUSE | Ferr_Suppressor |
| 13 | Gclc | GSH1_MOUSE | Ferr_Suppressor |
| 14 | Hspa8 | HSP7C_MOUSE | Ferr_Suppressor |
| 15 | Hspb1 | HSPB1_MOUSE | Ferr_Suppressor |
| 16 | Idh2 | IDHP_MOUSE | Ferr_Suppressor |
| 17 | Mt1 | MT1_MOUSE | Ferr_Suppressor |
| 18 | Mtor | MTOR_MOUSE | Ferr_Suppressor |
| 19 | Nfs1 | NFS1_MOUSE | Ferr_Suppressor |
| 20 | Nup62 | NUP62_MOUSE | Ferr_Suppressor |
| 21 | Otub1 | OTUB1_MOUSE | Ferr_Suppressor |
| 22 | Park7 | PARK7_MOUSE | Ferr_Suppressor |
| 23 | Pcbp1 | PCBP1_MOUSE | Ferr_Suppressor |
| 24 | Prdx1 | PRDX1_MOUSE | Ferr_Suppressor |
| 25 | Prdx6 | PRDX6_MOUSE | Ferr_Suppressor |
| 26 | Dhodh | PYRD_MOUSE | Ferr_Suppressor |
| 27 | Sqstm1 | SQSTM_MOUSE | Ferr_Suppressor |
| 28 | Tigar | TIGAR_MOUSE | Ferr_Suppressor |
| 29 | Trim11 | TRI11_MOUSE | Ferr_Suppressor |
| 30 | Txndc12 | TXD12_MOUSE | Ferr_Suppressor |
| 31 | Tmsb4x | TYB4_MOUSE | Ferr_Suppressor |

| | | | |
|---|---|---|---|
| 32 | Aco1 | ACOHC_MOUSE | Ferr_Driver |
| 33 | Acsf2 | ACSF2_MOUSE | Ferr_Driver |
| 34 | Atg5 | ATG5_MOUSE | Ferr_Driver |
| 35 | Cirbp | CIRBP_MOUSE | Ferr_Driver |
| 36 | Elavl1 | ELAV1_MOUSE | Ferr_Driver |
| 37 | Gsk3b | GSK3B_MOUSE | Ferr_Driver |
| 38 | Hmgb1 | HMGB1_MOUSE | Ferr_Driver |
| 39 | Idh1 | IDHC_MOUSE | Ferr_Driver |
| 40 | Mapk1 | MK01_MOUSE | Ferr_Driver |
| 41 | Mapk3 | MK03_MOUSE | Ferr_Driver |
| 42 | Pebp1 | PEBP1_MOUSE | Ferr_Driver |
| 43 | Pten | PTEN_MOUSE | Ferr_Driver |
| 44 | Fth1 | FRIH_MOUSE | Ferr_Marker |
| 45 | Gpx4 | GPX4_MOUSE | Ferr_Marker |
| 46 | Slc7a11 | XCT_MOUSE | Ferr_Marker |

**Table S5. Ferroptosis-associated proteins identified in A53T vs Control mice in the skeletal muscle.**

| S. No. | Genes | Gene_Name | Group |
|---|---|---|---|
| 1 | Got1 | AATC_MOUSE | Ferr_Suppressor |
| 2 | Aldh2 | ALDH2_MOUSE | Ferr_Suppressor |
| 3 | Hspa5 | BIP_MOUSE | Ferr_Suppressor |
| 4 | Cav1 | CAV1_MOUSE | Ferr_Suppressor |
| 5 | Cisd2 | CISD2_MOUSE | Ferr_Suppressor |
| 6 | Eno1 | ENOA_MOUSE | Ferr_Suppressor |
| 7 | Fxn | FRDA_MOUSE | Ferr_Suppressor |
| 8 | Ftl1 | FRIL1_MOUSE | Ferr_Suppressor |
| 9 | G3bp1 | G3BP1_MOUSE | Ferr_Suppressor |
| 10 | Hspa8 | HSP7C_MOUSE | Ferr_Suppressor |
| 11 | Hspb1 | HSPB1_MOUSE | Ferr_Suppressor |
| 12 | Idh2 | IDHP_MOUSE | Ferr_Suppressor |
| 13 | Mgst1 | MGST1_MOUSE | Ferr_Suppressor |
| 14 | Nfs1 | NFS1_MOUSE | Ferr_Suppressor |
| 15 | Otub1 | OTUB1_MOUSE | Ferr_Suppressor |
| 16 | Park7 | PARK7_MOUSE | Ferr_Suppressor |
| 17 | Pcbp1 | PCBP1_MOUSE | Ferr_Suppressor |
| 18 | Prdx1 | PRDX1_MOUSE | Ferr_Suppressor |

| | | | |
|---|---|---|---|
| 19 | Prdx6 | PRDX6_MOUSE | Ferr_Suppressor |
| 20 | Dhodh | PYRD_MOUSE | Ferr_Suppressor |
| 21 | Tigar | TIGAR_MOUSE | Ferr_Suppressor |
| 22 | Txndc12 | TXD12_MOUSE | Ferr_Suppressor |
| 23 | Tmsb4x | TYB4_MOUSE | Ferr_Suppressor |
| 24 | Prkaa2 | AAPK2_MOUSE | Ferr_Suppressor |
| 25 | Aco1 | ACOHC_MOUSE | Ferr_Driver |
| 26 | Cd36 | CD36_MOUSE | Ferr_Driver |
| 27 | Cirbp | CIRBP_MOUSE | Ferr_Driver |
| 28 | Fabp4 | FABP4_MOUSE | Ferr_Driver |
| 29 | Gsk3b | GSK3B_MOUSE | Ferr_Driver |
| 30 | Hmgb1 | HMGB1_MOUSE | Ferr_Driver |
| 31 | Idh1 | IDHC_MOUSE | Ferr_Driver |
| 32 | Mapk1 | MK01_MOUSE | Ferr_Driver |
| 33 | Mapk3 | MK03_MOUSE | Ferr_Driver |
| 34 | Pebp1 | PEBP1_MOUSE | Ferr_Driver |
| 35 | Ptbp1 | PTBP1_MOUSE | Ferr_Driver |
| 36 | Fth1 | FRIH_MOUSE | Ferr_Marker |
| 37 | Gpx4 | GPX4_MOUSE | Ferr_Marker |
| 38 | Tfrc | TFR1_MOUSE | Ferr_Marker |

**Table S6. Surface proteins identified in N2A cells upon αSyn-PFFs exposure.**

| S. No. | Gene | Fold Change | p-value |
|---|---|---|---|
| 1 | Vdac1 | 1.7289 | 0.0073854 |
| 2 | Tfrc | 5.3766 | 0.011404 |
| 3 | Rpn2 | 0.1931 | 0.034901 |
| 4 | Rpn1 | 1.8998 | 0.12252 |
| 5 | Eno1 | 0.50079 | 0.12718 |
| 6 | Slc3a2 | 0.74423 | 0.15718 |
| 7 | Ckap4 | 1.1303 | 0.17064 |
| 8 | Sidt2 | 0.39867 | 0.25334 |
| 9 | Emc1 | 0.36135 | 0.29227 |
| 10 | Nup210 | 1.4338 | 0.29667 |
| 11 | Atp1b3 | 1.4762 | 0.3005 |
| 12 | Atp1a1 | 1.8085 | 0.31787 |
| 13 | Itgb1 | 1.7322 | 0.39084 |
| 14 | Bsg | 1.1555 | 0.39789 |

| | | | |
|---|---|---|---|
| 15 | Ap2m1 | 1.5918 | 0.40831 |
| 16 | Plpp1 | 2.4806 | 0.4142 |
| 17 | Bst2 | 1.0675 | 0.45828 |
| 18 | Nrp2 | 1.4365 | 0.54244 |
| 19 | Calu | 2.0824 | 0.59405 |
| 20 | Emb | 1.081 | 0.67922 |
| 21 | Stim1 | 0.94825 | 0.69315 |
| 22 | Unc5c | 0.9039 | 0.7246 |
| 23 | Slc2a1 | 1.0115 | 0.7429 |
| 24 | Tmx3 | 1.1449 | 0.75733 |
| 25 | Havcr2 | 0.99312 | 0.77593 |
| 26 | H2-K1 | 1.6027 | 0.79369 |
| 27 | Hnrnpk | 1.0135 | 0.83781 |

**Table S7. Surface proteins identified in C2C12 cells upon αSyn-PFFs exposure.**

| S. No. | Gene | Fold change | p-value |
|---|---|---|---|
| 1 | Itgav | 0.17613 | 0.00043818 |
| 2 | Plpp1 | 3.1457 | 0.0037549 |
| 3 | Tfrc | 5.2046 | 0.0090746 |
| 4 | Ckap4 | 1.7402 | 0.015245 |
| 5 | Stim1 | 2.6139 | 0.023062 |
| 6 | Hnrnpk | 1.83 | 0.028466 |
| 7 | Ncam1 | 2.8435 | 0.030181 |
| 8 | Fbn1 | 0.080671 | 0.030965 |
| 9 | Lamp1 | 2.5132 | 0.033982 |
| 10 | Eno1 | 3.6768 | 0.044943 |
| 11 | Sidt2 | 2.0579 | 0.045418 |
| 12 | Rpn1 | 0.71163 | 0.05745 |
| 13 | Itgb1 | 2.882 | 0.085377 |
| 14 | Slc3a2 | 3.0258 | 0.096388 |
| 15 | Emc1 | 12.897 | 0.10069 |
| 16 | Bsg | 2.0193 | 0.10219 |
| 17 | Rpn2 | 1.362 | 0.13728 |
| 18 | Bgn | 1.9007 | 0.16781 |
| 19 | Fn1 | 1.8858 | 0.18175 |
| 20 | Ybx1 | 0.3261 | 0.20497 |
| 21 | Tmx3 | 1.3345 | 0.23243 |

| | | | |
|---|---|---|---|
| 22 | Fam171a2 | 1.7636 | 0.28003 |
| 23 | Vdac1 | 1.4646 | 0.32137 |
| 24 | Adgre5 | 0.58345 | 0.35884 |
| 25 | Itga7 | 1.4511 | 0.37082 |
| 26 | Slc2a1 | 0.78279 | 0.39745 |
| 27 | Cd63 | 1.1676 | 0.48514 |
| 28 | Atp1a1 | 1.2584 | 0.52075 |
| 29 | Slc6a6 | 0.80628 | 0.55617 |
| 30 | Lrp1 | 0.78922 | 0.60869 |
| 31 | Ncln | 0.54614 | 0.65112 |
| 32 | Cdh13 | 1.0274 | 0.65188 |
| 33 | Sgcd | 0.92445 | 0.6627 |
| 34 | Acp2 | 0.95282 | 0.66549 |
| 35 | Anxa2 | 0.86412 | 0.68801 |
| 36 | Cd80 | 0.15238 | 0.70193 |
| 37 | Cnnm4 | 1.4212 | 0.71805 |
| 38 | Cdh15 | 0.72823 | 0.72275 |
| 39 | Gpc1 | 1.0656 | 0.72494 |
| 40 | Ap2m1 | 1.0021 | 0.73598 |
| 41 | Psap | 1.1016 | 0.7641 |
| 42 | Ctsd | 1.0015 | 0.82748 |
| 43 | Atp1b3 | 0.98051 | 0.84076 |
| 44 | Ssr2 | 0.8919 | 0.8909 |
| 45 | Calu | 1.0663 | 0.95228 |
| 46 | Hspg2 | 1.5105 | 0.96615 |
| 47 | Itgb5 | 0.6463 | 0.96911 |
